# A comparative analysis of Parkinson’s disease and inflammatory bowel disease gut microbiomes highlights shared depletions in key butyrate-producing bacteria

**DOI:** 10.1101/2024.04.26.591350

**Authors:** Maeve E. Krueger, Jake Sondag Boles, Zachary D. Simon, Stephan D. Alvarez, Nikolaus R. McFarland, Michael S. Okun, Ellen M. Zimmermann, Christopher E. Forsmark, Malú Gámez Tansey

**Affiliations:** Department of Neuroscience, College of Medicine, University of Florida, Gainesville, FL, USA; Center for Translational Research in Neurodegenerative Disease, College of Medicine, University of Florida, Gainesville, FL, USA; McKnight Brain Institute, University of Florida, Gainesville, FL, USA; Department of Neurology, College of Medicine, University of Florida, Gainesville, FL, USA; Norman Fixel Institute for Neurological Diseases, University of Florida; Department of Medicine, Division of Gastroenterology, College of Medicine, University of Florida, Gainesville, FL, USA

## Abstract

Epidemiological studies reveal that a diagnosis of inflammatory bowel disease (IBD) is associated with an increased risk of developing Parkinson’s disease (PD). The presence of gut dysbiosis has been documented in both PD and IBD patients, however it is currently unknown how alterations in the gut microbiome may contribute to the epidemiological link between both diseases. To identify shared and distinct features of the PD and IBD microbiome, we performed the first joint analysis of 54 PD, 26 IBD, and 16 healthy control gut metagenomes recruited from clinics at the University of Florida, and directly compared the gut microbiomes from PD and IBD persons. Larger, publicly available PD and IBD metagenomic datasets were also analyzed to validate and extend our findings. Depletions in short-chain fatty acid (SCFA) producing bacteria, including *Roseburia intestinalis, Faecalibacterium prausnitzii, Anaerostipes hadrus,* and *Eubacterium rectale*, as well as depletions in SCFA synthesis pathways, were demonstrated across PD and IBD datasets. We posit that direct comparison of PD and IBD gut microbiomes will be important in identifying features within the IBD gut which may be associated with PD. The data revealed a consistent depletion in SCFA-producing bacteria across both PD and IBD, suggesting that loss of these microbes may influence the pathophysiology of both disease states.

## Introduction

Parkinson’s disease (PD) is a neurodegenerative disorder frequently associated with motor symptoms such bradykinesia, resting tremor, rigidity, and postural instability, PD is also associated with a host of non-motor symptoms many of which may possibly appear prior to motor symptom onset^1,2^. Accumulating evidence suggests the gut may play an important role in the pathogenesis of PD, with reports of up to 80% of patients experiencing gastrointestinal (GI) symptoms such as constipation^3^ and 20-30% reporting constipation in the prodromal phase^4,5^. Constipation has been associated with an increased risk for PD and continues to be listed among the most prevalent prodromal PD symptoms reported, appearing in some cases 10-20 years before diagnosis^6^. The appearance of constipation during the prodromal period may possibly reflect a body-first PD subtype. Some experts have opined that this body-type PD subtype is associated with autonomic dysfunction possibly resulting from abnormal α-synuclein (α-syn) aggregation in enteric neurons^7,8^ The theory proposes potential spread of pathology to the brain, with impairment of noradrenergic nuclei within the locus coeruleus. Interestingly, substantia nigra which is the compartment with dopamine cells becomes affected later in the process^4,9^. Increased levels of inflammatory markers seem to be asscociated with these changes and have been shown in animal models. These changes include IL-1α, IL-1β, CXCL8, C-reactive protein, and calprotectin in the PD gut. There has been a reported increase in intestinal permeability with experiments showing increased levels of α-1-antitrypsin and zonulin as well as aberrant distribution of tight junction proteins in human colon biopsies^10–14^.

An epidemiological association has been reported between inflammatory bowel disease (IBD) and PD, with IBD patients manifesting an increased risk of developing PD later in life^15–17^. IBD, which is characterized by the chronic relapsing of severe inflammation within the GI tract, serves as an umbrella diagnosis including Crohn’s disease (CD), which can affect many areas of the digestive tract, and ulcerative colitis (UC), which is more localized to the colon^18^. A recent comprehensive meta-analysis of 14 studies involving over 13.4 million patients concluded that IBD patients had a moderately increased risk (17%) of later developing PD^19^. A report by Peter et al. found this risk of developing PD was reduced by 78% if IBD patients were treated with TNF-inhibitors. This finding potentially suggested that chronic systemic inflammation was possibly important in the pathogenesis of both of these conditions^20^.

Emerging research highlights a compelling association between chronic inflammatory diseases and alterations in the composition of gut microbiota. Microbiome dysbiosis, which is characterized by disruptions in the balance of beneficial and pathogenic microbes, has been identified in patients with PD, IBD, and many other autoimmune and neurodegenerative diseases^21,22^. In PD, enrichments in pathogenic taxa such as *Escherichia coli* and *Klebsiella pneumoniae* have been reported^23^.

Depletions of beneficial, short-chain fatty acid (SCFA)-producing bacteria like *Roseburia intestinalis* and *Faecalibacterium prausnitizii* have also been detected.^23–26^ These findings seem to coincide with reduced concentrations of SCFAs in stool^11^. Acetate, propionate, and butyrate are known to be the most abundant SCFAs, predominately produced by anaerobic microbes in the gut via fermentation of dietary fibers^27^. These molecules thus serve a variety of potentially beneficial functions including providing direct fuel for colonocytes, increasing intestinal barrier integrity via the regulation of tight-junction proteins, suppressing inflammation by NFκB and MAPK signaling cascade inhibition, and enhancing the integrity of the blood-brain barrier^28,29^.

In IBD, the characterization of dysbiosis fluctuates between periods of active disease and remission^30^ and there is a manifestation of specific differences in dysbiosis between UC and CD diagnoses^31,32^. Overall, investigations of the IBD microbiome have reported depletions of *F. prausnitzii* and *R. intestinalis,* as well as a variety of other SCFA-producing *Clostridium* species^27,33,34^. Loss of these microbes seem to correspond to decreased concentrations of butyrate and acetate in the stool of IBD patients^33^. Enrichments of pathogenic *E. coli* have also been consistently reported in the IBD gut microbiome^35^, especially among CD patients. Finally, adherent invasive *E. coli* has been potentially implicated in the pathogenesis of the disease^36^.

While review of IBD and PD microbiome literature reveals similarities in the characterization of dysbiosis observed in both conditions, to our knowledge there has never been a direct comparison between PD and IBD gut microbiomes. Directly comparing the gut microbiomes of PD and IBD patients enables the exploration of a potential causal link between IBD and PD by the examination of microbial compositions and functional profiles. These insights could provide information to drive knowledge into the potential mechanisms linking gut microbiome alterations in IBD to increased risk of PD. To bridge this gap, we conducted a fecal metagenomic analysis of PD, IBD, and healthy control patients who were recruited from neurology and GI clinics at the University of Florida. We also analyzed larger, publicly available PD and IBD metagenomic datasets in an effort to enhance the robustness and generalizability of our findings, and to enable a deeper understanding of the shared features underlying PD and IBD. We demonstrate that in our sample that the IBD gut microbiome was altered, with a greater number of species both enriched and depleted when we compared directly to PD cases. In our study, IBD did not appear to share the enrichments in *Bifidobacterium* and *Lactobacillus* commonly reported to be associated with PD. Our data revealed a consistent depletion in SCFA-producing bacteria across both PD and IBD, suggesting that loss of these microbes may influence the pathophysiology of both diseases.

## Results

### UFPF subject demographics

As outlined in Table 1, our UFPF study consisted of 16 control, 26 IBD, and 54 PD subjects. As expected based on disease demographics, there was a significant difference in the average age between cohorts (*p* < 0.001). There were no significant differences in smoking history or estimated lifetime caffeine intake. 88.9% of PD subjects reported taking Carbidopa/Levodopa (*p* < 0.001) and 55.6% reported use of other PD medications (*p* < 0.001). 23.1% of IBD subjects reported use of anti-TNF medications (*p* = 0.001) and 80.8% reported other anti-inflammatory medication use (*p* < 0.001). Indigestion medication, depression/anxiety medication, and iron supplementation also differed significantly between groups (*p* = 0.015; *p* = 0.037, *p* = 0.006). Examination of potential associations between taxa and medication demonstrated that the use of anti-TNF medications was associated with enrichment of *Faecalimonas* in IBD (Supplementary Table 5).

**Table 1.**
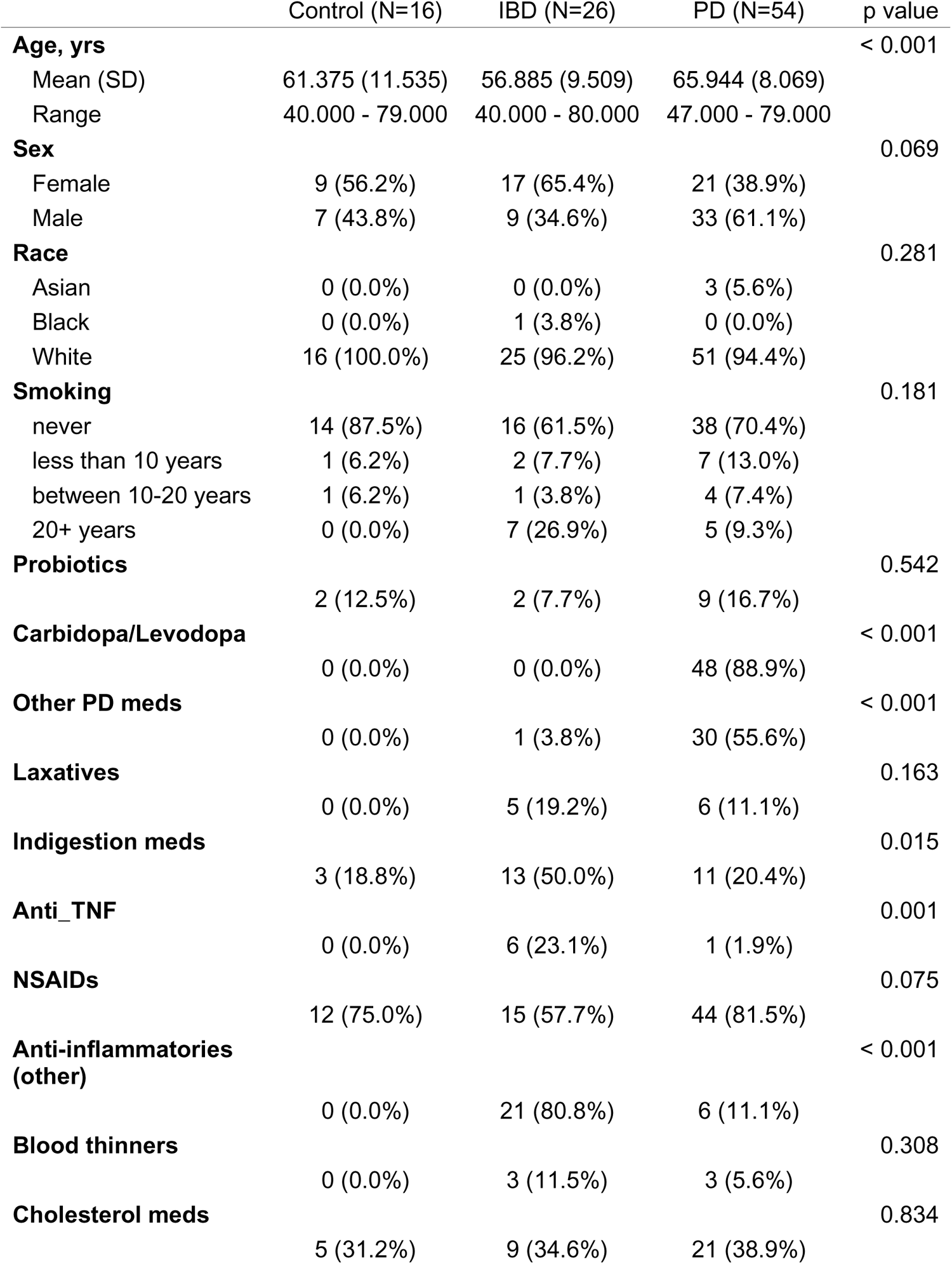
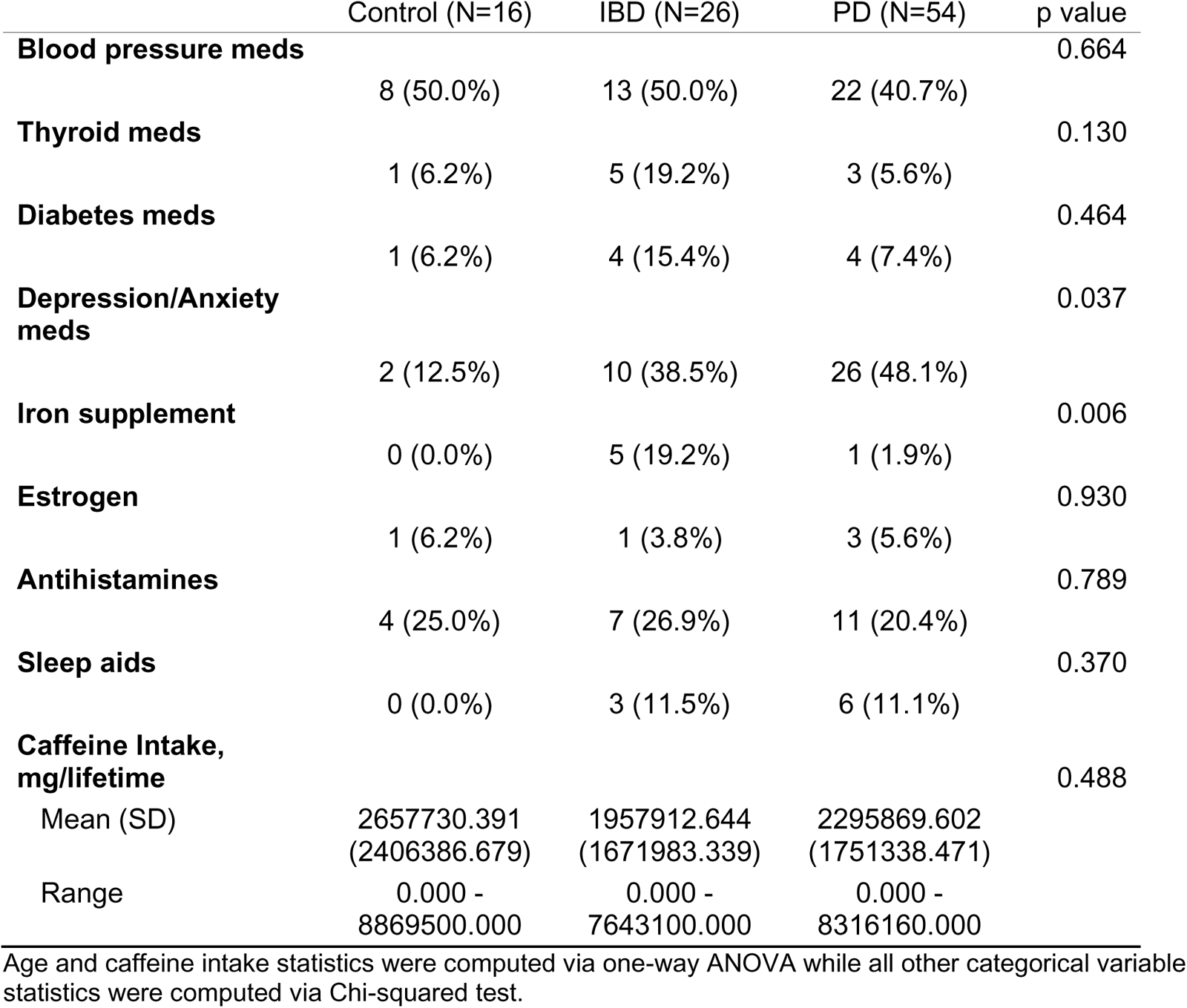
UFPF Subject Demographics and Metadata.

|  | Control (N=16) | IBD (N=26) | PD (N=54) | p value |
| --- | --- | --- | --- | --- |
| **Age, yrs** |  |  |  | < 0.001 |
| Mean (SD) | 61.375 (11.535) | 56.885 (9.509) | 65.944 (8.069) |  |
| Range | 40.000 - 79.000 | 40.000 - 80.000 | 47.000 - 79.000 |  |
| **Sex** |  |  |  | 0.069 |
| Female | 9 (56.2%) | 17 (65.4%) | 21 (38.9%) |  |
| Male | 7 (43.8%) | 9 (34.6%) | 33 (61.1%) |  |
| **Race** |  |  |  | 0.281 |
| Asian | 0 (0.0%) | 0 (0.0%) | 3 (5.6%) |  |
| Black | 0 (0.0%) | 1 (3.8%) | 0 (0.0%) |  |
| White | 16 (100.0%) | 25 (96.2%) | 51 (94.4%) |  |
| **Smoking** |  |  |  | 0.181 |
| never | 14 (87.5%) | 16 (61.5%) | 38 (70.4%) |  |
| less than 10 years | 1 (6.2%) | 2 (7.7%) | 7 (13.0%) |  |
| between 10-20 years | 1 (6.2%) | 1 (3.8%) | 4 (7.4%) |  |
| 20+ years | 0 (0.0%) | 7 (26.9%) | 5 (9.3%) |  |
| **Probiotics** |  |  |  | 0.542 |
|  | 2 (12.5%) | 2 (7.7%) | 9 (16.7%) |  |
| **Carbidopa/Levodopa** |  |  |  | < 0.001 |
|  | 0 (0.0%) | 0 (0.0%) | 48 (88.9%) |  |
| **Other PD meds** |  |  |  | < 0.001 |
|  | 0 (0.0%) | 1 (3.8%) | 30 (55.6%) |  |
| **Laxatives** |  |  |  | 0.163 |
|  | 0 (0.0%) | 5 (19.2%) | 6 (11.1%) |  |
| **Indigestion meds** |  |  |  | 0.015 |
|  | 3 (18.8%) | 13 (50.0%) | 11 (20.4%) |  |
| **Anti_TNF** |  |  |  | 0.001 |
|  | 0 (0.0%) | 6 (23.1%) | 1 (1.9%) |  |
| **NSAIDs** |  |  |  | 0.075 |
|  | 12 (75.0%) | 15 (57.7%) | 44 (81.5%) |  |
| **Anti-inflammatories (other)** |  |  |  | < 0.001 |
|  | 0 (0.0%) | 21 (80.8%) | 6 (11.1%) |  |
| **Blood thinners** |  |  |  | 0.308 |
|  | 0 (0.0%) | 3 (11.5%) | 3 (5.6%) |  |
| **Cholesterol meds** |  |  |  | 0.834 |
|  | 5 (31.2%) | 9 (34.6%) | 21 (38.9%) |  |
| **Blood pressure meds** |  |  |  | 0.664 |
|  | 8 (50.0%) | 13 (50.0%) | 22 (40.7%) |  |
| **Thyroid meds** |  |  |  | 0.130 |
|  | 1 (6.2%) | 5 (19.2%) | 3 (5.6%) |  |
| **Diabetes meds** |  |  |  | 0.464 |
|  | 1 (6.2%) | 4 (15.4%) | 4 (7.4%) |  |
| **Depression/Anxiety meds** |  |  |  | 0.037 |
|  | 2 (12.5%) | 10 (38.5%) | 26 (48.1%) |  |
| **Iron supplement** |  |  |  | 0.006 |
|  | 0 (0.0%) | 5 (19.2%) | 1 (1.9%) |  |
| **Estrogen** |  |  |  | 0.930 |
|  | 1 (6.2%) | 1 (3.8%) | 3 (5.6%) |  |
| **Antihistamines** |  |  |  | 0.789 |
|  | 4 (25.0%) | 7 (26.9%) | 11 (20.4%) |  |
| **Sleep aids** |  |  |  | 0.370 |
|  | 0 (0.0%) | 3 (11.5%) | 6 (11.1%) |  |
| **Caffeine Intake, mg/lifetime** |  |  |  | 0.488 |
| Mean (SD) | 2657730.391<br>(2406386.679) | 1957912.644<br>(1671983.339) | 2295869.602<br>(1751338.471) |  |
| Range | 0.000 -<br>8869500.000 | 0.000 -<br>7643100.000 | 0.000 -<br>8316160.000 |  |

Age and caffeine intake statistics were computed via one-way ANOVA while all other categorical variable statistics were computed via Chi-squared test.

### UFPF PD and IBD metagenomes demonstrate taxonomic and functional differences

In order to compare the gut microbiomes of our PD, IBD, and control patients, we performed taxonomic and functional profiling in addition to ordination analyses in order to assess the differences between the cohorts. In Figure 1A, a principal coordinate analysis using Aitchison distances demonstrated no difference between PD and control by PERMANOVA (*p* = 0.066) and PERMDISP (*p* = 0.496). However, IBD exhibited more localized clustering and dispersion that differed from both controls (PERMANOVA, *p* = 0.003; PERMDISP, *p* = 0.001) and PD (PERMANOVA, *p* = 0.003; PERMDISP, *p* = 0.001). To determine which taxa were driving the differences observed between cohorts, a differential abundance analysis was performed using ANCOM-BC2^37^. When performed at the species-level, only *Faecalimonas umbilicata* appeared significantly enriched in IBD compared to healthy control (Supplementary Table 1). When performed at the genus-level, two genera, *Klebsiella* and *Faecalimonas*, were significantly associated with IBD (Figure 1B). Both genera were significantly enriched in IBD compared to control and significantly depleted in PD when compared to IBD, but not compared to healthy controls. The full list of taxa with statistics from the genus-level ANCOM-BC2 analysis can be found in Supplementary Table 2.

**Figure 1:**
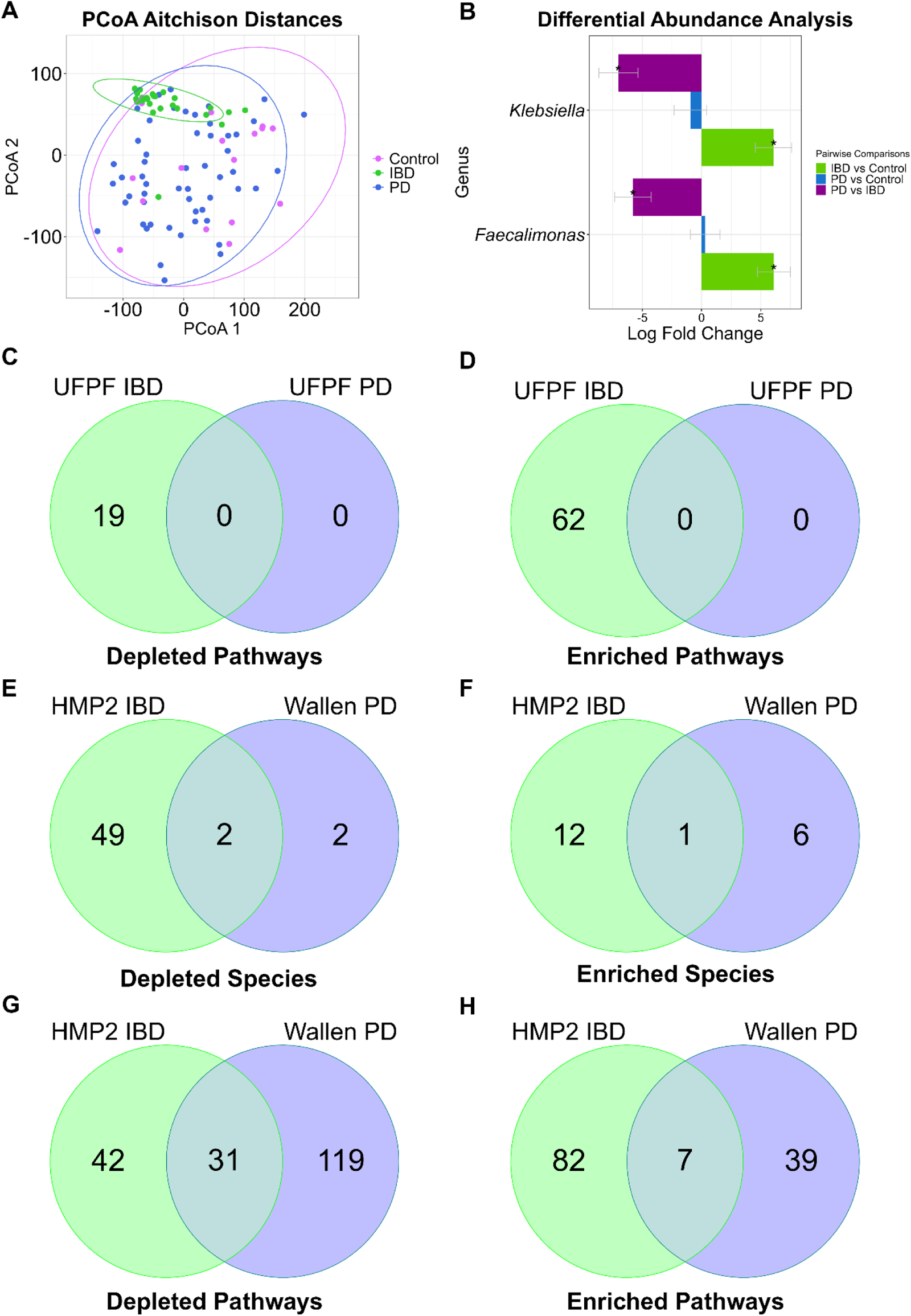
PCoA and ANCOM-BC2 analysis reveals taxonomic and functional differences between UFPF IBD and PD gut metagenomes while ANCOM-BC2 analysis of Wallen PD and HMP2 IBD metagenomes reveals enrichment and depletion of shared species and metabolic pathways. (A) Principal coordinate analysis (PCoA) using Aitchison distances between 54 PD (blue), 26 IBD (green), and 16 healthy control (pink) metagenomes with all species. (B) ANCOM-BC2 genus-level taxonomic differential abundance analysis reveals IBD-associated taxa, FDR < 0.05 with Benjamini Hochberg correction. Venn diagrams summarize the number of significant (*q-value* < 0.05) PD and IBD-associated MetaCyc pathways that were depleted and enriched (-/+ log fold change) in the UFPF dataset (C-D) and number of significant species (E-F) and MetaCyc pathways (G-H) depleted and enriched in the HMP2 IBD and Wallen PD datasets from ANCOM-BC2 differential abundance analysis.

Metagenomic analysis permits the evaluation of not only taxonomic differences, but also differences in functional potential, or the capacity of a microbial community to perform various biological functions based on the microbial genes present in each sample. In this study, we used HUMAnN3.5 to map gene families to functional units within the MetaCyc knowledgebase, allowing us to assess the abundance of specific metabolic pathways associated with the genes. Venn diagrams summarize the significant PD and IBD-associated MetaCyc pathways from ANCOM-BC2^37^ differential abundance analysis (Figure 1C-D). The significantly depleted and enriched pathways in IBD can be seen in Supplementary Table 4 while the full list of pathways can be found in Supplementary Table 3. No pathways were uniquely associated with PD (Figure 1C). The significantly depleted IBD-associated pathways demonstrate depletions in SCFA synthesis in relation to butyrate (CENTFERM-PWY; PWY-5676), as well as anaerobic energy metabolism (PWY-7383), gluconeogenesis (PWY66-399), and L-glutamate (PWY-5505) and L-methionine biosynthesis (HSERMETANA-PWY). Significantly enriched IBD-pathways show increases in amino acid degradation (ARGDEG-PWY; ORNARGDEG-PWY; AST-PWY; PWY0-461), polymyxin resistance (PWY0-1338), menaquinol biosynthesis (PWY-5862; PWY-5845; PWY-5860; PWY-5850; PWY-5896), and lipid-A biosynthesis (KDO-NAGLIPASYN-PWY) (Figure 1D).

### Wallen PD and HMP2 IBD subject demographics

In order to validate and expand upon the findings of our UFPF study, we reanalyzed larger, publicly availably PD and IBD datasets by repeating the same kind of ordination and differential abundance analyses in those datasets that we performed with the UFPF study data in order to compare the gut microbiomes of PD and control patients and IBD and non-IBD patients, respectively. The PD dataset, referred to as “Wallen PD”, utilized data from Wallen et al.^23^ downloaded via Zenodo [https://zenodo.org/doi/10.5281/zenodo.7246184] consisting of 490 PD and 234 neurologically healthy control samples (referred to as “control”). According to their report, the majority of control subjects in this cohort are spousal controls. There was a significant difference in sex (*p* < 0.001) and average age between PD and control (*p* < 0.001). 44.4% of PD subjects reported experiencing constipation (*p* < 0.001) while only 7.5% reported experiencing diarrhea compared to 12.6% of control subjects (*p =* 0.033). 31.4% PD subjects reported laxative use (*p* < 0.001). Pain medication, depression/anxiety/mood medication, birth control or estrogen, antihistamines, probiotics, and sleep aids also differed between groups (*p* = 0.03; *p* < 0.001, *p* < 0.001, *p* < 0.001, *p* = 0.011, *p* < 0.001). Detailed demographic information and metadata can be found in Supplementary Table 6.

The publicly available IBD dataset, referred to as “HMP2 IBD”, utilized data dated 2018-08-20 from the Human Microbiome Project 2 downloaded via https://www.ibdmdb.org/results. Due to the large age range of the HMP2 project, this dataset was filtered to include only subjects 40 years and older. This filtered dataset consisted of 198 IBD (101 CD, 97 UC) and 139 non-IBD samples collected from Cedars-Sinai and Massachusetts General Hospital. The HMP2 data did not collect spousal or healthy controls, but rather non-IBD subjects refer to subjects that were seen by the GI physician but were not diagnosed with IBD. There was a significant difference in sex between IBD and non-IBD (*p* < 0.001). 34.2% of IBD subjects reported probiotic use within the past 7 days (*p* < 0.001) while 10.6% of IBD subjects reported immunosuppressant use (*p* < 0.001) and 15.2% reported antibiotic use (*p* < 0.001). 47.9% of IBD subjects reported experiencing diarrhea in the past two-weeks (*p* < 0.001), and 31.2% reported a prior bowel surgery (*p* < 0.001). Detailed demographic information and metadata can be found in Supplementary Table 7. These PD and IBD datasets were statistically analyzed separately. The updated, computationally-intensive differential abundance analysis tool ANCOM-BC2^37^ was utilized to identify PD and IBD-associated taxa and pathways, which was not previously used in the published reports of these datasets as it had not yet been developed.

### Wallen PD and HMP2 IBD metagenomes display shared species and metabolic pathways

In the analysis of the Wallen PD dataset, PCoA revealed Wallen PD differed significantly from Control by both PERMANOVA (p = 0.001) and PERMDISP (p = 0.001) (Supplementary Figure 1A). Differential abundance analysis of the Wallen PD dataset revealed 11 species significantly associated with PD (Tables 2-3). Some of these were species associated with pathogenicity such as enrichments in *S. mutans, A. oris*, and *E. coli*. Other species enriched included *B. dentium, Eisenbergiella tayi, Ruminococcaceae bacterium D5*, and *C. leptum* (Table 2). Differential abundance analysis performed at the genus-level also demonstrated enriched *Lactobacillus* and *Enterococcus* which were not apparent at the species-level (Supplementary Table 8). Depleted taxa included *R. lactaris*, as well as *R. intestinalis, B. wexlerae*, and *A. hadrus*, reflecting a loss of prominent SCFA-producing bacteria in PD (Table 3).

**Table 2.** Wallen PD-Enriched Species.

| Species | Log-Fold Change | Standard Error | P-value | Adj. P-value |
| --- | --- | --- | --- | --- |
| *Eisenbergiella_tayi* | 1.40675355 | 0.44738066 | 0.00166415 | 0.03605665 |
| *Ruminococcaceae_bacterium_D5* | 1.56308293 | 0.42282987 | 0.0002184 | 0.00709814 |
| *Escherichia_coli* | 1.58740948 | 0.51939915 | 0.0022413 | 0.04370541 |
| *Clostridium_leptum* | 1.63026977 | 0.3979779 | 4.1966E-05 | 0.00204582 |
| *Streptococcus_mutans* | 1.88616654 | 0.41840646 | 6.5449E-06 | 0.00042542 |
| *Actinomyces_oris* | 2.07388674 | 0.41366513 | 5.3464E-07 | 0.00010426 |
| *Bifidobacterium_dentium* | 2.09821209 | 0.45715773 | 4.4391E-06 | 0.00042542 |

**Table 3.** Wallen PD-Depleted Species.

| Species | Log-Fold Change | Standard Error | P-value | Adj. P-value |
| --- | --- | --- | --- | --- |
| *Roseburia_intestinalis* | -1.91171 | 0.512205 | 0.00019 | 0.007098 |
| *Blautia_wexlerae* | -1.4315 | 0.418764 | 0.00063 | 0.017547 |
| *Ruminococcus_lactaris* | -1.62883 | 0.488916 | 0.000864 | 0.021054 |
| *Anaerostipes_hadrus* | -1.51515 | 0.506813 | 0.002794 | 0.049526 |

In the analysis of the HMP2 IBD dataset, PCoA revealed IBD differed significantly from non-IBD by PERMANOVA (p = 0.001) but not by PERMDISP (p = 0.832) (Supplementary Figure 1B). Performing differential abundance analysis of the HMP2 IBD dataset revealed 64 IBD-associated species (Tables 4-5). 13 of these 64 taxa were detectably enriched including *E. coli*, several *Clostridium* species, *Ruminococcus gnavus*, and others (Table 4). At the genus-level, enrichment in taxa that did not reach significance at the species-level such as *Propionibacteriaceae, Eggerthella, and Klebsiella* were also detected (Supplementary Table 9). Depletions in 51 species were evident, reflecting an IBD microbiome that had lost numerous beneficial bacteria including *Bifidobacterium adolescentis, A. muciniphila, P. copri,* and prominent SCFA-producers like *R. instestinalis, R. hominis, E. rectale, and F. prausnitizii* (Table 5).

**Table 4.**
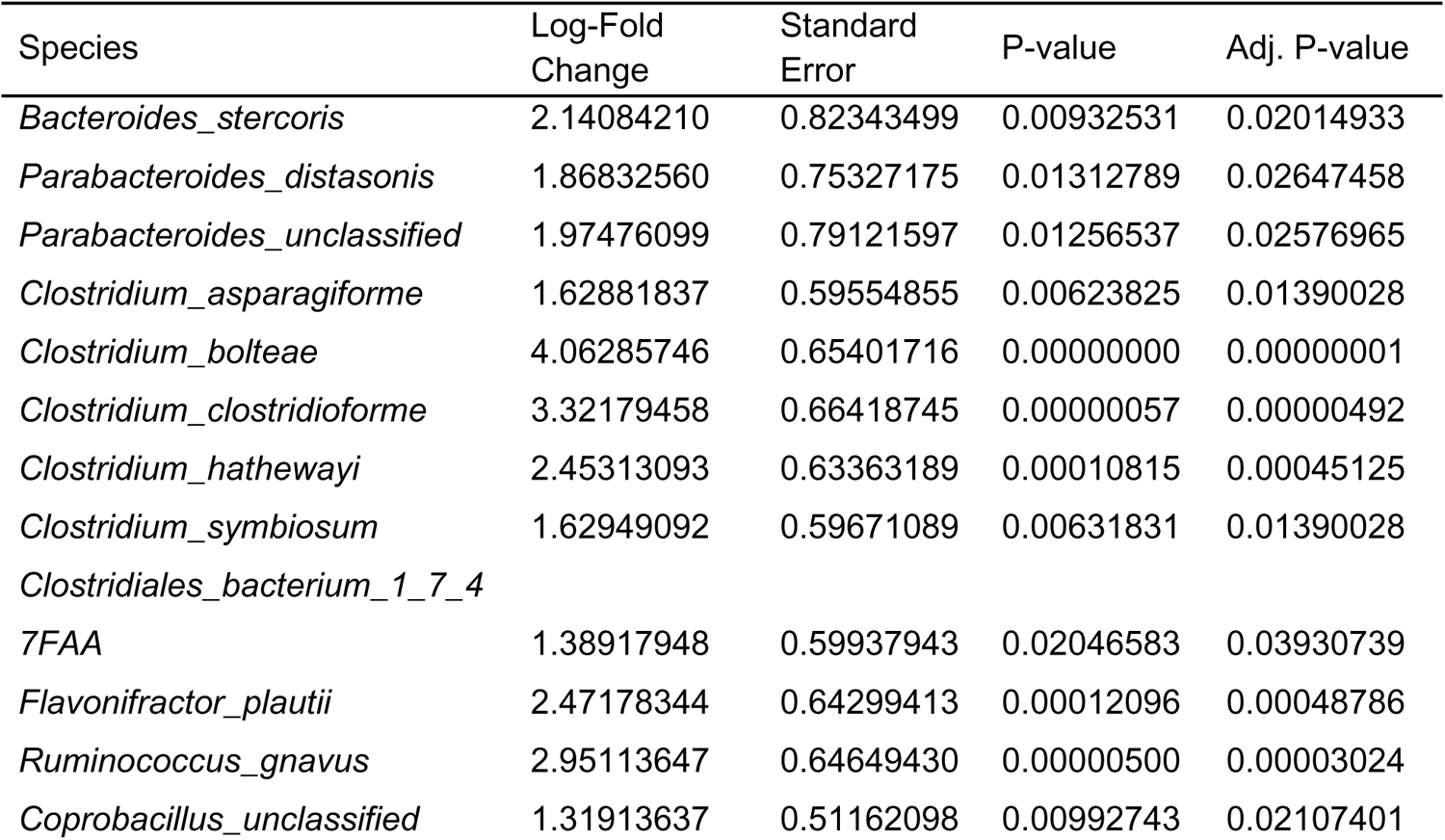
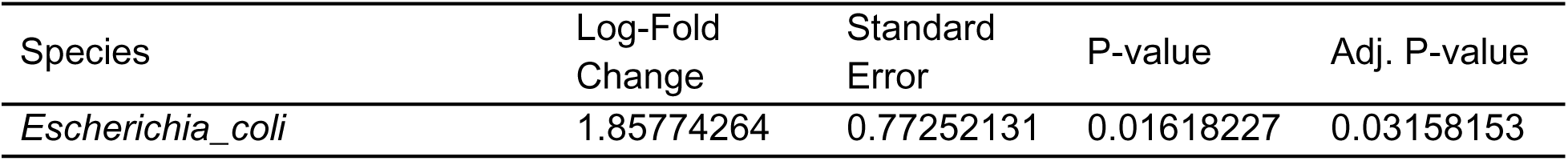
HMP2 IBD-Enriched Species.

**Table 5.**
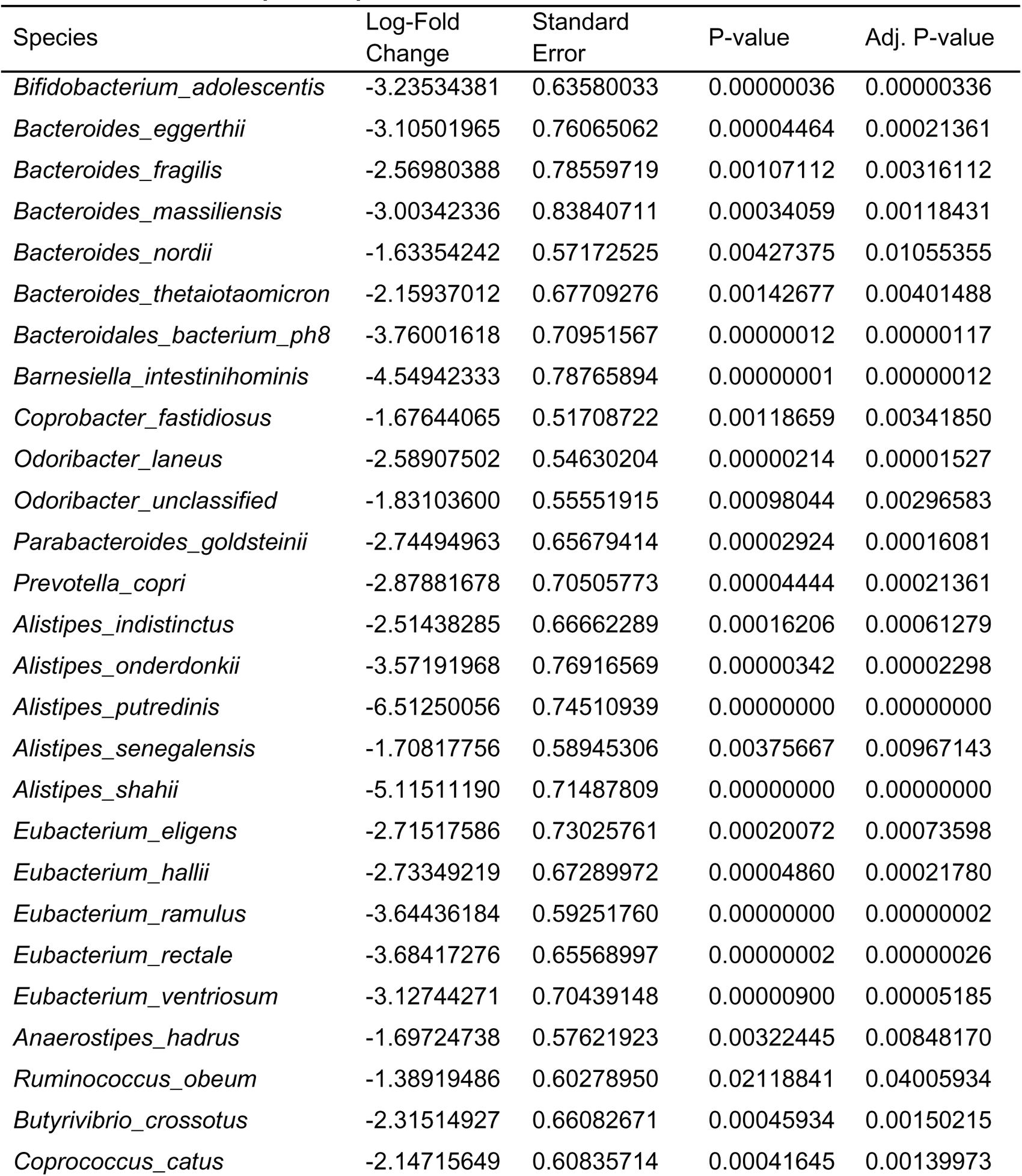
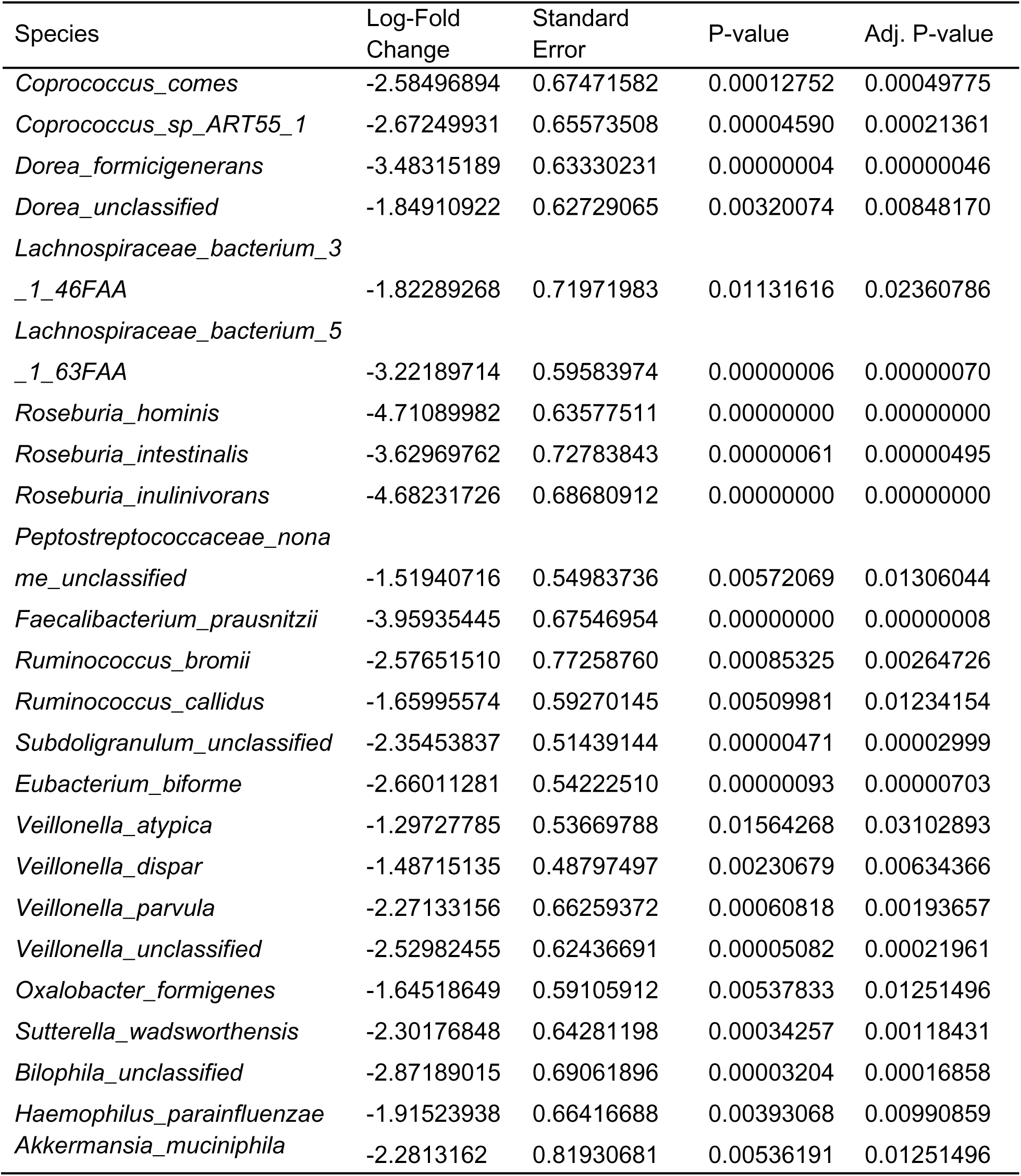
HMP2 IBD-Depleted Species.

Although we have not directly compared the Wallen PD and HMP2 IBD datasets in an integrated analysis due to substantial differences between them in subject demographics, geographic location, sample collection, and processing methods, we have compared the results from the differential abundance analyses of the Wallen PD and HMP2 IBD datasets, outlining the features that were significantly depleted and enriched in each. Two SCFA-producing species were significantly depleted in both the Wallen PD and HMP2 IBD cohorts, *R. intestinalis* and *A. hadrus* (Figure 1E), while *E. coli* was the only commonly enriched species between both datasets (Figure 1F).

Following functional analysis, Wallen PD displayed depletion in 150 MetaCyc pathways (Figure 1G; Supplementary Table 10). Some of these depleted pathways included depletion in SCFA synthesis (PWY-7111; PWY-5676; PWY-5100), L-glutamate/L-glutamine biosynthesis (PWY-5505), tryptophan biosynthesis (PWY-6629; TRPSYN-PWY), glycogen biosynthesis (GLYCOGENSYNTH-PWY), L-arginine biosynthesis (PWY-5154; PWY-7400; ARGSYN-PWY), peptidoglycan synthesis (PEPTIDOGLYCANSYN-PWY; PWY0-1586; PWY-6385; PWY-6471), and lactic acid fermentation (ANAEROFRUCAT-PWY). HMP2 IBD displayed depletion in 73 pathways (Figure 1G; Supplementary Table 11), including depletions seen in SCFA synthesis pathways (PWY-5100; CENTFERM-PWY; P108-PWY; PWY-5676), L-methionine biosynthesis (HSERMETANA-PWY; PWY-5345; PWY-5347; METSYN-PWY; HOMOSER-METSYN-PWY), and other amino acid synthesis (PWY-3001; P4-PWY; SER-GLYSYN-PWY; PWY-5104; PWY-5103; DAPLYSINESYN-PWY). Both Wallen PD and HMP2 IBD shared depletions in 31 pathways including glycogen biosynthesis (GLYCOGENSYNTH-PWY), L-methionine biosynthesis (HSERMETANA-PWY; MET-SAM-PWY; PWY-5347), butanoate II production (PWY-5676), acetate and lactate II production (PWY-5100), and L-serine, glycine, and L-isoleucine biosynthesis (PWY-5103; PWY-5104; SER-GLYSYN-PWY) (Table 6).

**Table 6.** Wallen + HMP2 Shared-Depleted Pathways.

| Pathway | LFC IBD | LFC PD | SE IBD | SE PD | P-val IBD | P-val PD | Adj. P-val IBD | Adj. P-val PD |
| --- | --- | --- | --- | --- | --- | --- | --- | --- |
| ARO-PWY: chorismate biosynthesis I | -0.29148 | -0.27773 | 0.122171 | 0.052381 | 0.017041 | 1.14E-07 | 0.032766 | 8.67E-07 |
| CALVIN-PWY: Calvin-Benson-Bassham cycle | -0.32084 | -0.28289 | 0.119924 | 0.052958 | 0.007465 | 9.20E-08 | 0.016137 | 7.28E-07 |
| COMPLETE-ARO-PWY: superpathway of aromatic amino acid biosynthesis | -0.26981 | -0.28087 | 0.120141 | 0.05228 | 0.024718 | 7.77E-08 | 0.045137 | 6.92E-07 |
| GALACT-GLUCUROCAT-PWY: superpathway of hexuronide and hexuronate degradation | -0.87943 | -0.39838 | 0.18829 | 0.070417 | 3.00E-06 | 1.54E-08 | 2.39E-05 | 1.71E-07 |
| GALACTUROCAT-PWY: D-galacturonate degradation I | -0.8727 | -0.355 | 0.178319 | 0.065544 | 9.88E-07 | 6.09E-08 | 1.04E-05 | 5.70E-07 |
| GLUCONEO-PWY: gluconeogenesis I | -1.40898 | -0.1672 | 0.267139 | 0.058327 | 1.33E-07 | 0.004149 | 1.87E-06 | 0.009233 |
| GLYCOGENSYNTH-PWY: glycogen biosynthesis I (from ADP-D-Glucose) | -0.66634 | -0.34797 | 0.125074 | 0.05708 | 9.96E-08 | 1.09E-09 | 1.54E-06 | 1.76E-08 |
| HSERMETANA-PWY: L-methionine biosynthesis III | -1.27925 | -0.26788 | 0.144284 | 0.054501 | 7.57E-19 | 8.88E-07 | 2.22E-16 | 4.05E-06 |
| MET-SAM-PWY: superpathway of S-adenosyl-L-methionine biosynthesis | -0.91905 | -0.13744 | 0.17217 | 0.04387 | 9.40E-08 | 0.001731 | 1.54E-06 | 0.00434 |
| OANTIGEN-PWY: O-antigen building blocks biosynthesis (E. coli) | -0.75381 | -0.14447 | 0.195226 | 0.058941 | 0.000113 | 0.014241 | 0.000377 | 0.027856 |
| POLYAMSYN-PWY: superpathway of polyamine biosynthesis I | -0.38197 | -0.25849 | 0.167875 | 0.088222 | 0.022888 | 0.00339 | 0.042056 | 0.007887 |

**Table 6. continued**
| Pathway | LFC IBD | LFC PD | SE IBD | SE PD | P-val IBD | P-val PD | Adj. P-val IBD | Adj. P-val PD |
| --- | --- | --- | --- | --- | --- | --- | --- | --- |
| PWY-3001: superpathway of L-isoleucine biosynthesis I | -0.37103 | -0.24752 | 0.129881 | 0.053178 | 0.004281 | 3.25E-06 | 0.00991 | 1.28E-05 |
| PWY-5030: L-histidine degradation III | -0.90291 | -0.26229 | 0.182239 | 0.073915 | 7.25E-07 | 0.000387 | 8.20E-06 | 0.001143 |
| PWY-5100: pyruvate fermentation to acetate and lactate II | -0.57761 | -0.23385 | 0.107856 | 0.057603 | 8.54E-08 | 4.91E-05 | 1.54E-06 | 0.00016 |
| PWY-5103: L-isoleucine biosynthesis III | -0.64729 | -0.26055 | 0.114225 | 0.053268 | 1.45E-08 | 1.00E-06 | 3.29E-07 | 4.51E-06 |
| PWY-5104: L-isoleucine biosynthesis IV | -0.8076 | -0.28291 | 0.177297 | 0.072693 | 5.24E-06 | 9.95E-05 | 3.58E-05 | 0.000319 |
| PWY-5188: tetrapyrrole biosynthesis I (from glutamate) | -0.33798 | -0.31542 | 0.125421 | 0.061306 | 0.007044 | 2.68E-07 | 0.01557 | 1.67E-06 |
| PWY-5347: superpathway of L-methionine biosynthesis (transsulfuration) | -0.87051 | -0.14811 | 0.163216 | 0.041941 | 9.64E-08 | 0.000414 | 1.54E-06 | 0.001207 |
| PWY-5676: acetyl-CoA fermentation to butanoate II | -0.67808 | -0.29927 | 0.25541 | 0.071867 | 0.007934 | 3.12E-05 | 0.016942 | 0.000104 |
| PWY-6121: 5-aminoimidazole ribonucleotide biosynthesis I | -0.39672 | -0.26551 | 0.121371 | 0.051842 | 0.001081 | 3.03E-07 | 0.002812 | 1.79E-06 |
| PWY-6122: 5-aminoimidazole ribonucleotide biosynthesis II | -0.39753 | -0.26462 | 0.12382 | 0.05199 | 0.001325 | 3.58E-07 | 0.003273 | 1.99E-06 |
| PWY-6277: superpathway of 5-aminoimidazole ribonucleotide biosynthesis | -0.39753 | -0.26462 | 0.12382 | 0.05199 | 0.001325 | 3.58E-07 | 0.003273 | 1.99E-06 |

**Table 6. continued**
| Pathway | LFC IBD | LFC PD | SE IBD | SE PD | P-val IBD | P-val PD | Adj. P-val IBD | Adj. P-val PD |
| --- | --- | --- | --- | --- | --- | --- | --- | --- |
| PWY-6507: 4-deoxy-L-threo-hex-4-enopyranuronate degradation | -0.87359 | -0.34376 | 0.199124 | 0.068093 | 1.15E-05 | 4.46E-07 | 6.89E-05 | 2.37E-06 |
| PWY-6527: stachyose degradation | -0.58024 | -0.41508 | 0.092017 | 0.059162 | 2.87E-10 | 2.28E-12 | 2.56E-08 | 1.35E-10 |
| PWY-6606: guanosine nucleotides degradation II | -0.7247 | -0.29096 | 0.145599 | 0.072241 | 6.45E-07 | 5.63E-05 | 7.58E-06 | 0.000182 |
| PWY-6737: starch degradation V | -0.4676 | -0.29141 | 0.134439 | 0.053123 | 0.000505 | 4.12E-08 | 0.001387 | 3.97E-07 |
| PWY-7199: pyrimidine deoxyribonucleosides salvage | -0.43523 | -0.26737 | 0.126335 | 0.053588 | 0.000571 | 6.06E-07 | 0.00154 | 2.95E-06 |
| PWY-7242: D-fructuronate degradation | -0.6141 | -0.42541 | 0.170857 | 0.069437 | 0.000325 | 8.98E-10 | 0.000947 | 1.68E-08 |
| SER-GLYSYN-PWY: superpathway of L-serine and glycine biosynthesis I | -0.52327 | -0.21743 | 0.121416 | 0.053556 | 1.63E-05 | 4.91E-05 | 9.10E-05 | 0.00016 |
| TRNA-CHARGING-PWY: tRNA charging | -0.43814 | -0.24706 | 0.097426 | 0.051629 | 6.89E-06 | 1.71E-06 | 4.50E-05 | 7.41E-06 |
| TRPSYN-PWY: L-tryptophan biosynthesis | -0.39722 | -0.29497 | 0.156557 | 0.055696 | 0.011173 | 1.18E-07 | 0.0225 | 8.78E-07 |

Wallen PD displayed enrichments in 46 pathways (Figure 1H; Supplementary Table 12). These pathways included L-arginine degradation (ARGDEG-PWY; ORNARGDEG-PWY), vitamin K2 synthesis (PWY-5838; PWY-5845; PWY-5897; PWY-5898; PWY-5899), L-threonine metabolism (THREOCAT-PWY), and other energy metabolism-related pathways. HMP2 IBD displayed enrichments in 89 pathways (Figure 1H; Supplementary Table 13). These pathways included L-arginine degradation (AST-PWY), antimicrobial resistance (PWY0-1338), lipid A biosynthesis (KDO-NAGLIPASYN-PWY), vitamin K2 biosynthesis (PWY-5845; PWY-5850; PWY-5896; PWY-5838), phospholipid biosynthesis (PHOSLIPSYN-PWY), heme biosynthesis (HEME-BIOSYNTHESIS-II; PWY-5918), and L-glutamate/glutamine biosynthesis (PWY-5505). Wallen PD and HMP2 IBD shared seven commonly enriched pathways including glucose degradation (GLUCOSE1PMETAB-PWY), ketogluconate metabolism (KETOGLUCONMET-PWY), trehalose degradation (PWY-2723), vitamin K2 biosynthesis (PWY-5838; PWY-5845; PWY-5862), pyrimidine deoxyribonucleotides *de novo* biosynthesis (PWY-7198) (Table 7).

**Table 7.** Wallen + HMP2 Shared-Enriched Pathways.

| Pathway | LFC IBD | LFC PD | SE IBD | SE PD | P-val IBD | P-val PD | Adj. P-val IBD | Adj. P-val PD |
| --- | --- | --- | --- | --- | --- | --- | --- | --- |
| GLUCOSE1PMETAB-PWY: glucose and glucose-1-phosphate degradation | 0.845569 | 0.579749 | 0.236682 | 0.177505 | 0.000353 | 0.00109 | 0.001009 | 0.002881 |
| KETOGLUCONMET-PWY: ketogluconate metabolism | 0.768626 | 0.54102 | 0.192885 | 0.212445 | 6.75E-05 | 0.010877 | 0.000254 | 0.021753 |
| PWY-2723: trehalose degradation V | 0.824011 | 0.583506 | 0.191237 | 0.184198 | 1.64E-05 | 0.001536 | 9.10E-05 | 0.003934 |
| PWY-5838: superpathway of menaquinol-8 biosynthesis I | 0.94766 | 0.468934 | 0.245903 | 0.178944 | 0.000116 | 0.008779 | 0.000384 | 0.018065 |
| PWY-5845: superpathway of menaquinol-9 biosynthesis | 0.941523 | 0.621274 | 0.237433 | 0.203845 | 7.33E-05 | 0.002305 | 0.000263 | 0.00566 |
| PWY-5862: superpathway of demethylmenaquinol-9 biosynthesis | 0.917845 | 0.580048 | 0.225414 | 0.1944 | 4.66E-05 | 0.002847 | 0.00019 | 0.006848 |
| PWY-7198: pyrimidine deoxyribonucleotides de novo biosynthesis IV | 0.61013 | 0.119364 | 0.248912 | 0.041559 | 0.014239 | 0.004077 | 0.028285 | 0.009128 |

### Module analysis reveals similar abundances of key taxa across PD and IBD cohorts

In order to relate the UFPF study findings to the significant features found in the Wallen PD and HMP2 IBD datasets, we created disease-associated taxonomic modules based on these significant features and evaluated their abundances in our UFPF cohort. Significantly enriched and depleted species from the ANCOM-BC2 differential abundance analysis of the Wallen and HMP2 datasets were extracted into separate feature lists. Module scores were then calculated, comprised of the average abundance of species found within those feature lists in a dataset of interest, such as within the UFPF PD, IBD, and control cohorts.

UFPF IBD displayed a significantly increased abundance of species found within the Enriched in Wallen PD module, but we did not detect an enrichment of these species in UFPF PD (Figure 2A). However, UFPF PD displayed a significantly decreased abundance of species found within the Depleted in Wallen PD module compared to control (Figure 2B). UFPF IBD displayed a significantly increased abundance of species found within the Enriched in HMP2 IBD module compared to UFPF PD and control (Figure 2C), and both UFPF IBD and PD displayed significantly decreased abundances of species found within the Depleted in HMP2 IBD module (Figure 2D). Notably, this included SCFA producers *F. prausnitzii, E. rectale, R. intestinalis, A. hadrus,* and others.

**Figure 2.**
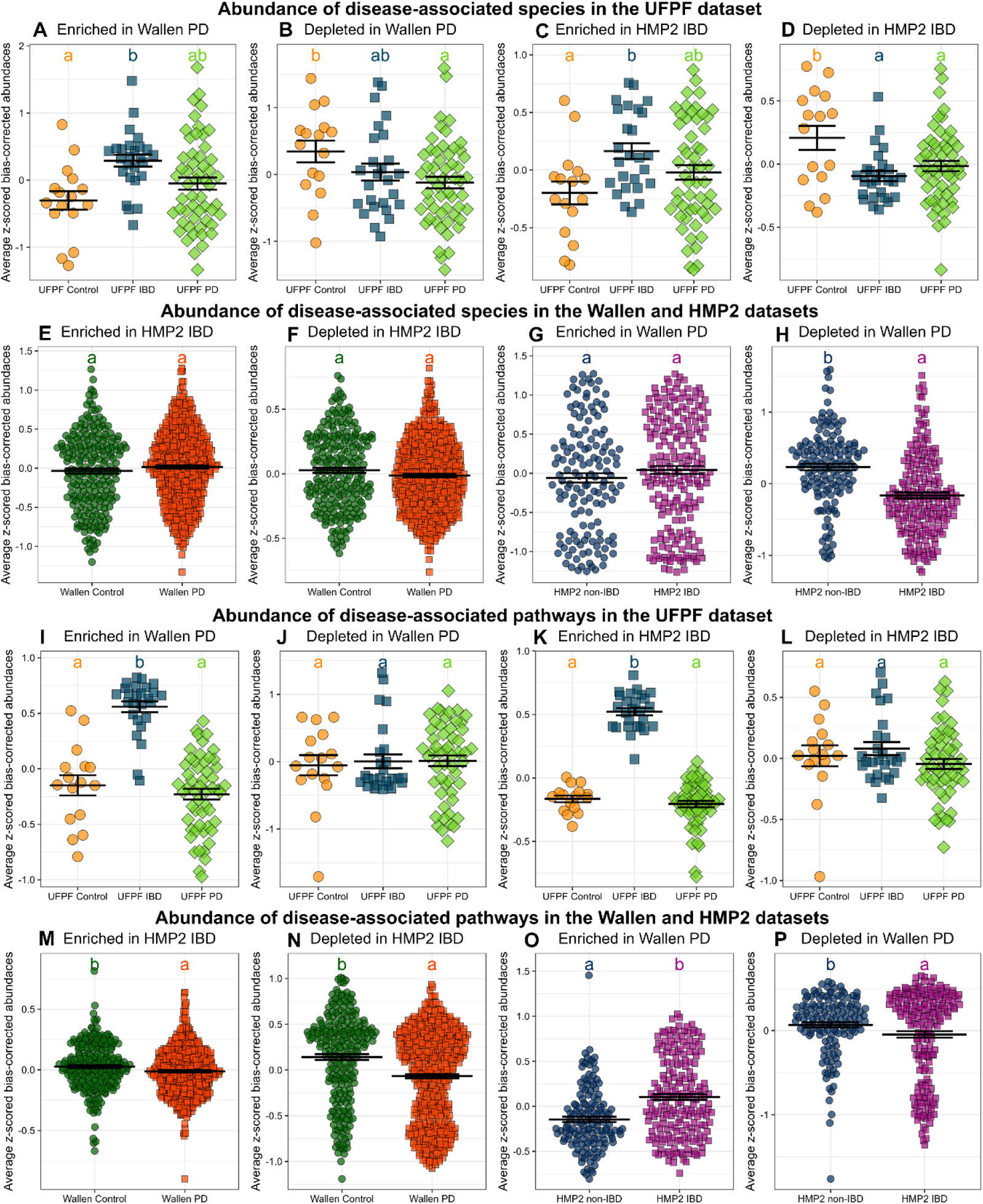
Species module analysis reveals depletion of shared species across PD and IBD while pathway module analysis reveals variability of depleted and enriched pathways across PD and IBD. (A-D) The abundance of significant species from the Wallen PD and HMP2 IBD datasets within the UFPF PD, IBD, and control cohorts are plotted. Modules are denoted by the title on each individual plot. Significantly enriched and depleted species were extracted into separate feature lists from the ANCOM-BC2 differential abundance analysis of the Wallen and HMP2 datasets. Calculated module scores were comprised of the average abundance of those species found within those feature lists in the UFPF PD, IBD, and control cohorts, using the R WGCNA package. One-way ANOVAs were performed followed by calculation of estimated marginal means and pairwise comparisons. Compact letter display was used to display pairwise comparisons, as letters that are different from each other are significantly different (*p* < 0.05). (E-F) The abundance of significant species from the HMP2 IBD dataset within the Wallen PD dataset are plotted and (G-H) the abundance of significant species from the Wallen PD dataset within the HMP2 IBD dataset are plotted. The same statistical methods as described above were used, but the module scores were comprised of the average abundance of species found within the feature lists in the HMP2 IBD and Wallen PD datasets. (I-L) The abundance of significant MetaCyc pathways from the Wallen PD and HMP2 IBD datasets within the UFPF PD, IBD, and control cohorts are plotted. Significantly enriched and depleted MetaCyc pathways were extracted into separate feature lists from the ANCOM-BC2 differential abundance analysis of the Wallen and HMP2 datasets. Calculated module scores were comprised of the average abundance of those pathways found within those feature lists in the UFPF PD, IBD, and control cohorts. The same statistical methods were used as described for (A-H). (M-N) The abundance of significant pathways from the HMP2 IBD dataset within the Wallen PD dataset are plotted and (O-P) the abundance of significant pathways from the Wallen PD dataset within the HMP2 IBD dataset are plotted. The same methods as described above were used, but the module scores were comprised of the average abundance of pathways found within the feature lists in the HMP2 IBD and Wallen PD datasets.

We employed the same strategy to compare the two publicly available datasets, examining the abundance of taxa identified in these disease-associated taxonomic modules in either dataset. Wallen PD did not display alterations in the abundance of species within the Enriched or Depleted in HMP2 IBD modules (Figure 2E-F). HMP2 IBD displayed no difference in the abundance of species found within the Enriched in Wallen PD module compared to non-IBD (Figure 2G). However, HMP2 IBD did display a significantly decreased abundance of species found within the Depleted in Wallen PD module (Figure 2H). Overall, UFPF PD and IBD cohorts shared similarities with larger Wallen PD and HMP2 IBD datasets, and there appeared to be similarities in shared depleted microbes across all PD and IBD datasets examined.

### Module analysis does not reveal consistent patterns in metabolic pathway abundance across PD and IBD

To compare functional alterations in the IBD and PD microbiomes, we grouped disease-associated MetaCyc pathways from the differential abundance analyses of the Wallen PD and HMP2 IBD datasets (Supplementary Tables 10-13) into co-abundance modules using the same methods as described above for the taxonomic module analysis. The UFPF IBD cohort displayed an increased abundance of pathways found within the Enriched in Wallen PD module compared to UFPF PD and control (Figure 2I). Interestingly, UFPF PD did not appear to share an increased abundance of pathways seen in this module. Further, there were no differences in the abundance of pathways found within the Depleted in Wallen PD module across UFPF IBD and PD compared to control (Figure 2J). UFPF IBD but not UFPF PD displayed a significantly increased abundance of pathways found within the Enriched in HMP2 IBD module relative to control (Figure 2K). Neither UFPF PD or IBD displayed altered abundances of pathways found within the Depleted in HMP2 IBD module (Figure 2L). Wallen PD displayed a significantly decreased abundance of pathways found within both the Enriched in HMP2 IBD module (Figure 2M) and the Depleted in HMP2 IBD module (Figure 2N) compared to control. Meanwhile, HMP2 IBD displayed a significantly increased abundance of pathways found within the Enriched in Wallen PD module compared to non-IBD (Figure 2O) as well as a significantly decreased abundance of pathways found within the Depleted in Wallen PD module (Figure 2P). Overall, results of the functional module analysis revealed the IBD cohorts consistently demonstrated increased abundances of pathways seen enriched in the Wallen PD dataset but did not appear to display further consistent patterns that were shared between PD and IBD.

## Discussion

Although several published studies have examined IBD and PD gut metagenomes, tour study was unique as it directly compared the gut metagenomes from IBD, PD, and control subjects. While some differences in the groups could not be controlled, such as difference in subject age and medication use, the control of the methodological variables was a strength of this study. Additionally, our analysis of the larger PD and IBD datasets utilizing updated bioinformatic tools which could be compared to previously published analyses allowed us to validate and expand upon the findings from our UFPF study dataset. Our study addressed a critical knowledge gap, and expanded beyond the epidemiological association between IBD and PD. We raise the hypothesis that the IBD patients who later develop PD may possibly exhibit similar metagenomic profiles to PD patients or may harbor unique features that elevate the risk for PD. These ideas can be fleshed out in future studies.

Our analysis revealed an enrichment of microbes associated with pathogenicity such as *E. coli, S. mutans, and A. oris* in Wallen PD, which was also seen in the original report by Wallen *et al.*^23^. Although these taxa were not significantly enriched in UFPF PD, increased abundance of *E. coli* is commonly associated with PD, with a previous report demonstrating *E. coli* was associated with worsened postural instability and gait difficulty in PD^38^. HMP2 IBD also displayed significant enrichment in *E. coli* as well as *Klebsiella*, at the genus-level,another pathogenic, gram-negative bacterium. Our UFPF IBD cohort also demonstrated significant enrichment in *Klebsiella.* Accompanying these taxonomic findings, our functional analyses demonstrated that both IBD datasets were uniquely associated with increased lipid A biosynthesis, a glycolipid that serves as a critical component of lipopolysaccharide (LPS)^23^ found in the outer membrane of gram-negative bacteria. While lipid A biosynthesis was significantly increased in PD in the original report by Wallen et al.^23^, it did not reach significance in our analysis.

Enrichments in members of the *Proteobacteria* phylum, such as *E. coli* and *Klebsiella pneumoniae*, are typically reported in the gut microbiomes of PD^23^ and IBD patients^39,40^ and have been implicated in the development of IBD itself^41^. Studies in murine models have demonstrated that targeted suppression of potential IBD-associated pathogens, *K. pneumoniae* and adherent invasive *E. coli* (AIEC), dampens intestinal inflammation and ameliorates colitis^42,43^. Experimental evidence suggests that gut inflammation in IBD may induce low-grade neuroinflammation, as murine models of colitis have also demonstrated evidence of BBB dysfunction^44^ as well as increased inflammatory cytokine levels, microglial activation, and loss of myelin^45–47^. Moreover, IBD patients have been shown to experience higher rates of depression which has been suggested to be a result of this low-grade neuroinflammation which is induced by the invasion of systemic immune factors and subsequent microglial activation in the brain^48^. This suggests that IBD patients exhibiting pronounced intestinal inflammation may be predisposed to heightened neuroinflammation, potentially amplifying the risk of developing PD.

Additionally, the bacterial amyloid produced by *E. coli*, curli, has been reported to induce α-syn misfolding in the gut and brain of α-syn-overexpressing (ASO) mice^49^. Curli has also been shown to promote α-syn aggregation inside neurons through a cross-seeding mechanism which revealed human α-syn-expressing *Caenorhabditis elegans*^50^ in a human sample study. This misfolded α-syn may occur within luminal-facing enteroendocrine cells (EECs) which express α-syn in the neuropod portion of the cell and are innervated by vagal afferents, making this an attractive hypothesis to possibly explain how α-syn misfolding in the gut could propagate to the brain via the vagus nerve in PD^51–53^. Given evidence that the gut environment can influence neuroinflammation and neuropathology via the gut-brain-axis, this idea we opine will warrant future investigation as to whether targeting pathogenic microbes, like *E. coli* or *K. pneumoniae*, in the gut of IBD patients could reduce PD risk.

Given the marked enrichment of *Faecalimonas* observed in our UFPF IBD cohort, which was comprised of patients in remission, this association holds particular relevance for IBD patients, as the use of anti-TNF therapies may possibly induce positive shifts in gut microbiome composition that could potentially be protective against the development of future inflammation-associated diseases like PD. These findings may thus provide insight into a potential gut microbiome-derived mechanism that could contribute to the understanding as to why there is a 78% reduction in the incidence of PD among IBD patients who had taken anti-TNF therapies as reported by Peter *et al.*^20^.

*A. muciniphila*, a mucin-degrading microbe largely thought to be protective against inflammation, is often enriched in neuroinflammatory conditions including Alzheimer’s disease and multiple sclerosis^57,58^. While increases in *Akkermansia* have previously been associated with PD^24,59,60^, the IBD microbiome consistently exhibits depletion, potentially due to the loss of the intestinal outer mucus layer in active IBD, which this microbe utilizes as a fuel source^61,62^. Enrichment of *A. muciniphila* did not reach significance by differential abundance analysis in UFPF PD or Wallen PD, but HMP2 IBD displayed a significant decrease in *A. muciniphila*. The lack of *A. muciniphila* depletion in the UFPF IBD cohort may reflect the remission status of the patients, as Vigsnæs *et al*. demonstrated that UC patients in remission showed no significant difference *in A. muciniphila* while UC patients with active disease showed a significant loss of *A. muciniphila*^63^. While there are ongoing investigations exploring *A. muciniphila* administration as a therapeutic, not all evidence suggests this microbe is beneficial.

Germ-free IL-10 KO mice administered multiple doses of *A. muciniphila* can spontaneously develop colitis^64^. Additionally, *A. muciniphila* has been associated with the development of some neuropsychiatric conditions^65^. Given this conflicting evidence and because the role of *A. muciniphila* in PD and IBD remains elusive, further research should be directed to explore the mechanisms underlying its involvement and therapeutic implications in both PD and IBD .

Notably, investigations into the PD microbiome consistently report significant enrichments in commensal species of *Bifidobacterium* and *Lactobacillus*^23,24,26^. It is not known why these microbes are enriched in PD, but one hypothesis suggests it may be the result of a compensatory response to developing dysbiosis^23^. Enrichment in *Lactobacillus* and *B. dentium* were demonstrated in Wallen PD. *B. dentium* was not detected in the IBD cohorts examined, however, previous reports have demonstrated depletion of *Bifidobacterium* species in the IBD microbiome. While more evidence in the IBD literature exists for other *Bifidobacterium* species such as *B. breve*, *B. longum*, and *B.infantis*^66–68^, one report suggests that *B. dentium* administration may aid in the enhancement of gut mucosal function by increasing mucin production^69^.

While numerous studies identify *B. dentium* as a probiotic flora, other reports speculate that it may contribute to gait issues in PD via this microbe’s ability to convert glutamate to GABA^70^, as elevated GABA levels in one study have been associated with a worsened PD gait^71^. Previous reports in PD patients have implicated overactive glutamate receptors in the promotion of neurodegeneration^72^ as well as identified increased glutamate degradation and low glutamate production in the PD gut metagenome^73^. While we did not detect increased glutamate degradation in the PD gut, the Wallen PD cohort did reveal reduced L-glutamate biosynthesis. The UFPF IBD cohort also displayed depletion in L-glutamate biosynthesis while the HMP2 IBD cohort displayed enrichment in this pathway. It is unclear why the IBD cohorts examined revealed different features, but loss of glutamate in the gut due to increased consumption or reduced production may have potentially contributed to intestinal inflammation as decreased glutamate has been suggested to contribute to mucosal damage^73^. Additional research could be directed to investigate the extent to which elevation in *Bifidobacterium* and *Lactobacillus* is a maladaptive or compensatory response in PD, the answer to which could impact the desirability and tolerability of supplementation with these bacteria in patients with PD.

The most striking similarity between IBD and PD across every dataset examined was a shared depletion in similar taxa, as demonstrated by our taxonomic module analysis (Figure 2). Many of these taxa are prominent SCFA-producing microbes including *A. hadrus, R. intestinalis*, *F. prausnitzii, B. wexlerae, E. rectale, E. hallii (*reclassified as *Anaerobutyricum hallii,)* and *E. eligens.* More specifically, a number of these microbes are major butyrate-producers in the colon including *R. intestinalis*, *F. prausnitzii, A. hadrus*, *A. hallii, and Eubacterium* species including *E. rectale and E. eligens*^74–79^. *F. prausnitzii,* a microbe known for its anti-inflammatory properties, has been shown in a 2,4,6-trinitrobenzene sulfonic acid (TNBS) colitis model to protect against colitis, improve gut dysbiosis, and decrease serum levels of TNF^79^. *B. wexlerae* administration has been shown to increase acetate and lactate levels, which can be used to produce butyrate via cross-feeding and promote the growth of other SCFA-producers^80^. Decreases in *B. wexlerae* have been associated with increased inflammation and insulin resistance^81^. Numerous publications have identified decreased abundance of SCFAs in IBD, especially butyrate, and their producers^27,33,82–84^ as well as in PD^11,29,78,85^. In our analysis, we also observed depletions in pathways involved in the synthesis of acetate and butyrate across PD and IBD datasets that accompany this loss of SCFA-producing bacteria, raising the interesting possibility that this loss may have possibly contributed to both disease processes.

SCFAs have been shown to exert a variety of immunomodulatory effects as they can bind to receptors on a variety of cell types, including myeloid cells, affording them the ability to influence epithelial immunity and immune tolerance^86^. Our group previously identified epigenetic signatures in immune cells from individuals with PD, notably neutrophils and monocytes, that were associated with butyrate levels in stool^87^. These epigenetic changes most closely resembled those we also found in IBD, both UC and CD, when compared to other neurological diseases like Alzheimer’s disease and bipolar disorder. This observation could provide further support for a unique association between IBD and PD, as opposed to other neurologic conditions, and could possibly implicate gut microbiota as the potential trigger for these epigenetic modifications.

Additional evidence points to butyrate potentially providing protection against neurodegeneration via the ability to inhibit histone deacetylates (HDAC)^88–91^. HDAC inhibitors have been shown to have neuroprotective effects in PD, and other neurodegenerative diseases, by increasing protein acetylation and promoting downstream gene expression and modulation of immune responses ^92–94^. In models of colitis, HDAC inhibitors have been shown to reduce colitis severity and decrease colonic TNF^95^. Notably, our data demonstrated a loss of major butyrate producers across the PD and IBD cohorts, leading us to posit that the loss of certain butyrate-producing bacteria in IBD may produce alterations in host immunity, fostering a gut microenvironment with a propensity to escalate inflammation. Additionally, a leaky gut characteristic of IBD could in turn potentially further promote neuroinflammation and heightened PD risk. It is tempting to speculate that loss of butyrate, and its function as an immune modulator and HDAC inhibitor, could contribute to the development or progression of both IBD and PD. This possibility raises the question that restoring the colonization of butyrate-producing microbes or restoring levels of butyrate itself in IBD may afford protection against or delay development of PD. In support of this idea, previous work from our group revealed that among subjects with PD, those with higher levels of butyrate in the stool reported a later age-at-onset of disease, suggesting a potentially protective effect of butyrate^11^. The effects of SCFA administration in PD patients have not been thoroughly explored; however this approach could be pursued in a clinical trial targeting early PD.

Additional findings from out data included shared enrichments in pathways related to Vitamin K2 biosynthesis seen in Wallen PD and HMP2 IBD cohorts. Menaquinone, Vitamin K2, can be synthesized by lactic acid bacteria in the gut and used by the host for a variety of functions including calcium metabolism and blood clotting^96^. Vitamin K2 is commonly depleted in IBD, which is likely a result of increased bleeding, anemia, and inflammation associated with active IBD^97^. Given the enrichment of these pathways in both PD and IBD, increases in Vitamin K2 synthesis could be due to underlying gut inflammation, potentially reflecting a dysregulated and complex microbe-host interaction. More targeted experiments will be required to parse out the functional significance of PD gut dysbiosis and what it may mean in terms of disease context and subsequent changes to a host immune response.

Wallen PD displayed enrichments in L-arginine degradation and depletions in L-arginine synthesis, which have been reported previously by Jo *et al*. in a study of the PD gut metagenome^73^. Both of the IBD cohorts examined here also demonstrated enrichment of L-arginine degradation pathways. L-arginine aids in inflammation regulation and can be oxidized to nitric oxide, and has been shown to play a beneficial role in neurogenesis, neuroplasticity, learning and memory^98^. Additionally, Wallen PD displayed decreased L-tryptophan biosynthesis while HMP2 IBD displayed enrichment in this pathway. It is well documented that PD patients display serotoninergic deficits^99^, and previous reports of the PD metagenome have identified increased consumption of tryptophan in PD^100,101^. Increases in tryptophan metabolism have been suggested to follow inflammation or infection and have been associated with IBD^102^. Therefore, we posit that it is possible that decreased tryptophan synthesis in both PD and IBD metagenomes could contribute to the serotonin dysregulation reported in these patients^99,103^. Additional studies will be required to confirm or refute this theory.

It is unclear whether the microbiome alterations observed in PD and IBD are a cause or consequence of disease. Given the complex interactions between the host immune system and microbiota in the gut, it is possible that inflammation promotes a dysbiotic microbiome, favoring the persistence of pathogenic microbes that can tolerate more extreme conditions. One study that administered 5% dextran sodium sulfate (DSS) in the drinking water of C57BL/6 mice for 5 days demonstrated decreased gut microbial diversity and dysbiosis characterized by enrichments in *Helicobacter* and *Escherichia*, which are reportedly enriched in IBD patients^104^. However, antibiotic treatment has been shown to ameliorate DSS-induced colitis^105^. In PD, a study using antibiotic-treated and germ-free α-syn overexpressing (ASO) mice demonstrated that gut microbiota were required for the development of motor deficits and robust α-syn pathology, while recolonization with fecal microbiota transplants (FMTs) from PD patients exacerbated motor impairments^106^. Given these findings, alterations to the microbiota or the presence of certain disease-specific microbes may be necessary to produce the immunologic dysregulation and pathology observed in IBD and PD. Future studies involving colonization of germ-free mice with FMTs from IBD patients would help establish the extent of colitogenicity conferred by the IBD gut microbiota and its effects on α-syn pathology in humanized α-syn mice.

There were several limitations to our study. We were limited in our analysis by the small and uneven sample sizes of our UFPF PD, IBD, and control cohorts. Given that the Wallen PD and HMP2 IBD datasets were unrelated, we did not statistically analyze them together which prevented us from running direct comparisons between the larger PD and IBD cohorts. Confirmatory studies with larger sample sizes of PD and IBD patients will be required in the future to firmly establish and further explore the gut microbiome as a potential link between these diseases. The UFPF PD and IBD cohorts did display some compositional differences compared to previously collected PD and IBD cohorts which may have reflected the inherent heterogeneity associated with microbiome investigations influenced by stool collection method, geographical location, diet, or other factors. Dietary information was not collected as part of the UFPF study, so we could not examine potential dietary influences on the gut microbiome alterations reported herein. It will be important for future analyses to examine the influence of dietary intake on the comparison of PD and IBD gut microbiomes. Additionally, the lack of an IBD-household control group limited our ability to examine disease-specific effects in the UFPF IBD cohort. UFPF IBD patients also partially ingested the colonoscopy prep polyethylene glycol just prior to stool sample collection. While it is known that gut microbiome composition is immediately altered the day after consumption of bowel preparation with polyethylene glycol, the UFPF IBD samples were collected upon first bowel movement after polyethylene glycol ingestion. However, it is still possible that a small amount of laxative influenced their microbiome composition prior to complete bowel emptying. It will be important for future studies to examine the acute effects of polyethylene glycol ingestion on the gut microbiome to determine if this significantly influences composition.

Despite the inherent variability observed between disease cohorts, it is clear that the UFPF PD and IBD cohorts share key similarities with the Wallen PD and HMP2 IBD cohorts respectively. When comparing PD and IBD microbiomes, depletion in major SCFA-producers, especially butyrate-producing bacteria, along with subsequent depletion in SCFA synthesis pathways, are notable features shared between the two diseases. The results of the present study should be followed up in larger PD and IBD cohorts in order to resolve variability encountered with the small sample sizes in the UFPF study and the limitations of the utilization of larger datasets from separate studies. Larger cohorts would also allow stratification of IBD into UC and CD, which could provide useful insights into unique PD risk depending on IBD diagnosis, especially given that differences have been reported between UC and CD gut microbiomes^31^. Due to the epidemiological association between IBD and PD, it could be hypothesized that the gut microbiomes of IBD patients that go on to develop PD may have similar metagenomic profiles to PD patients or contain features that uniquely increase the risk for PD. It is also possible that loss of butyrate-producing bacteria in IBD may produce alterations in host immunity and compromise peripheral immune tolerance, establishing a gut microenvironment with a propensity to promote systemic inflammation that could in turn lead to downstream neuroinflammation and increased PD risk. Incorporating metabolomic analyses into future investigations of PD and IBD gut microbiomes may thus be important in moving beyond SCFAs and in exploring microbial metabolites that could offer deeper insights into the functional consequences of microbial dysbiosis across these disease states. Metabolomic profiling has the potential to elucidate the specific microbial metabolites altered in PD and IBD, and may possibly reveal novel therapeutic targets that could modulate the increased risk of PD in IBD patients.

In summary, our findings on the existence of shared features of gut microbiome between IBD and PD especially in the characteristic of depletion of SCFA-producing bacteria could lay some of the foundations for future investigations into the mechanisms underlying these similarities. Notably, our findings hereby represent an advance for the field beyond a mere epidemiological association between IBD and PD risk.

## Methods

We have complied with all relevant ethical regulations and were approved by the Institutional Review Board (IRB) at the University of Florida. All subjects gave signed informed consent, approved by UF IRB.

Our study was conducted at the University of Florida by a team consisting of neurology and gastroenterology investigators who participated in study design and established subject enrollment procedures across University of Florida clinics. PD patients were recruited from the University of Florida Neuromedicine Clinic at the Fixel Institute for Neurodegenerative Diseases. IBD patients preparing for routine colonoscopy were recruited from the UF Gastrointestinal (GI) Clinic. Healthy control patients (referred to as “control”) were all PD-spousal controls with the exception of one subject that was an IBD patient-spousal control.

56 PD patients, 26 IBD patients, and 16 control patients were recruited between August 2020 and May 2023. Inclusion criteria for participation in this cross-sectional study included age 40-80 years, diagnosis of PD (based on MDS criteria^107^), IBD, or without either condition in the case of the control cohort. Exclusion criteria included PD individuals with active infections or active autoimmune or chronic inflammatory conditions (except for IBD) that require treatments with immunosuppressants, pregnancy, history of a blood transfusion within 4-weeks of recruitment, body weight less than 110 lbs., subjects who were perceived by the investigator to be unable to comply with study procedures, and treatment with immunosuppressants or antibiotics within one month of recruitment. Notably, all IBD subjects recruited for this study were declared in remission, as confirmed by colonic biopsies that were collected by the gastroenterologists as part of routine colonoscopies. With each potential participant, informed consent was conducted in a private setting where the research coordinator explained the study and the subject’s involvement thoroughly. Time was given for the subject to read in the informed consent form and ask any questions. If the subject elected to participate in the study, they signed the consent.

Subject questionnaires were used to collect medical history, medications, family history, alcohol and tobacco use, and caffeine intake. Caffeine intake was reported by subjects as average cups per day for “x” number of years of coffee, tea, and soda. Estimates for caffeine content of these three beverages were as follows: coffee (80mg), tea (40mg), and soda (25mg). PD subjects were also administered UPDRS, MOCA, Environmental Questionnaire and Medical History, and Schwab and England ADL questionnaire. Subject questionnaires utilized in this study can be found in the supplementary materials. Dietary information was not collected as part of this study.

### Stool Swab Collection

Study subjects were provided mail-in stool collection kits consisting of a BD BBL CultureSwab EZ II (Fisher Scientific, Catalog No. BD220145) along with instructions for stool sample collection (Supplementary Materials). Subjects were instructed to insert collection swab directly into the stool, or if not possible, rub the swab on stool residue present on toilet paper after wiping. Collection kits were mailed overnight and were placed in a freezer at -80°C upon receipt by study staff. IBD patients recruited were already undergoing routine colonoscopy, thus patients took the prep polyethylene glycol an osmotic laxative and were instructed to collect from the first bowel movement following ingestion of the prep. PD patients collected stool swabs without the influence of polyethylene glycol.

### DNA Extraction

For DNA extraction and sequencing, samples were sent on dry ice to CosmosID Inc. (Germantown, MD, USA). DNA was isolated using the QIAGEN DNeasy PowerSoil Pro Kit, according to the manufacturer’s protocol. DNA samples were quantified using the GloMax Plate Reader System (Promega) using the QuantiFluor dsDNA system (Promega) chemistry.

### Library Preparation and Sequencing

DNA libraries were prepared using the Nextera XT DNA Library Preparation Kit (Illumina) and IDT Unique Dual Indexes with total DNA input of 1ng. Genomic DNA was fragmented using a proportional amount of Illumina Nextera XT fragmentation enzyme. Unique dual indexes were added to each sample followed by 12 cycles of PCR to construct libraries. DNA libraries were purified using AMpure magnetic Beads (Beckman Coulter) and eluted in QIAGEN EB buffer. DNA libraries were quantified using Qubit 4 fluorometer and Qubit dsDNA HS Assay Kit. Libraries were then sequenced on Illumina NovaSeq 6000 platform 2x150bp.

### QC

All QC processing of sequence reads, as well as downstream profiling using MetaPhlAn4^108^ and HUMAnN3.5^109^, were performed in HiperGator, the University of Florida’s high-performance computing (HPC) cluster that permits intensive computational tasks. QC on paired-end sequences received from CosmosID Inc. were performed in accordance with the steps and parameters outlined in Wallen et al.^23^. All sequences that mapped to a human reference genome GRCh38.p13 with accession number GCA_000001405.28 [https://www.ncbi.nlm.nih.gov/assembly/GCA_000001405.28] were removed using BBSplit with default settings [https://sourceforge.net/projects/bbmap/]. The full code we used for performing QC can be found in the project’s GitHub page [https://github.com/maevekrueger/UFPF_metagenomics] under “Bioinformatic Processing of Sequences.” The resulting sequences consisted of 139,618 – 10,077,039 reads per sample. One PD sample had a depth of 26,329 reads and was subsequently excluded from all downstream analysis, as well as another PD sample which lacked any available metadata needed for analysis, leaving us with 54 PD samples available for this analysis.

### Taxonomic Profiling Using MetaPhlAn4

Following QC, sequences were profiled using MetaPhlAn4 [https://huttenhower.sph.harvard.edu/metaphlan] (RRID:SCR_004915) a computational tool permitting the taxonomic classification of microbial communities including bacteria, archaea, and eukaryotes^108^. MetaPhlAn4 relies on a robust database containing ∼5.1M unique clade-specific marker genes that allow for species and strain-level identification. Overall methods and settings for running MetaPhlAn4 were performed in accordance with those outlined in Wallen et al.^23^. The ChocoPhlAn database version used was mpa_vJun23_CHOCOPhlAnSGB_202307 [http://cmprod1.cibio.unitn.it/biobakery4/metaphlan_databases/]. Relative abundances were generated from MetaPhlAn4 using default settings. MetaPhlAn4 was run a second time with the ‘–unclassified-estimation’ flag added to generate relative abundance values that included the estimated abundance of microbes not found within the database. These values were used to calculate raw counts by multiplying the relative abundance values by the total read count of that sample. These counts were used in downstream analysis involving Aitchison distances and differential abundance analysis using ANCOM-BC2^37^. Individual sample files were merged using ‘merge_metaphlan_tables.py.’

### Functional Profiling Using HUMAnN3.5

Following QC, sequences were profiled for functional MetaCyc pathways using HUMAnN3.5^109^ [https://huttenhower.sph.harvard.edu/humann] (RRID:SCR_014620) to default parameters. Individual output files were merged using the ‘humann_join_tables’ command to achieve distinct output files: path_abundance.tsv and path_coverage.tsv.

### Analysis In R

Data wrangling and visualization were performed in R (version 4.2.3). For Venn diagram creation, the ggVennDiagram package was used [https://cran.r-project.org/web/packages/ggVennDiagram/index.html] was used. To create the metadata and demographics table, the arsenal package [https://cran.r-project.org/web/packages/arsenal/index.html] was used with the tableby() function. All code used to generate tables and figures for this paper is open access and available on this project’s GitHub page [https://github.com/maevekrueger/UFPF_metagenomics].

### PCoA

An Aitchison distance matrix was created by calculating Euclidean distances on centered log-ratio (CLR) transformed species-level counts. A small pseudo-count was added to counts before performing CLR transformation using the clr() function from the Compositions package [https://cran.r-project.org/web/packages/compositions/index.html]. A Euclidean distance matrix was then created using the vegdist() function, method “euclidean” from the vegan package [https://cran.r-project.org/web/packages/vegan/index.html] (RRID:SCR_011950).

Because we created a Euclidean distance matrix from CLR-transformed counts, this is now referred to as an Aitchison distance matrix. A principal coordinate analysis (PCoA) was performed on the Aitchison distance matrix using the pcoa() function from the ape package [https://cran.r-project.org/web/packages/ape/index.html] (RRID:SCR_017343). Eigenvalues and eigenvectors were extracted from the PCoA results and used in the creation of a scatter plot displaying samples in a two-dimensional space using ggplot2 [https://cran.r-project.org/web/packages/ggplot2/index.html] (RRID:SCR_014601).

PERMANOVA tests were conducted using adonis() from the vegan package and pairwise comparisons were conducted when appropriate, using the pairwiseAdonis package [https://github.com/pmartinezarbizu/pairwiseAdonis] with the Bonferroni’s *p*-value adjustment method. While PERMANOVA can indicate differences between groups, this may indicate differences in dispersion and/or location. PERMDISP is a multivariate extension of Levene’s test, permitting the evaluation of dispersion differences (variances) between groups of samples. Also from the vegan package, PERMDISP was performed using the betadisper() function and pairwise differences were assessed using permutest() on the results of betadisper().

### Differential Abundance Analysis with ANCOM-BC2

#### ANCOM-BC2

[https://www.bioconductor.org/packages/release/bioc/vignettes/ANCOMBC/inst/doc/AN COMBC2.html] (RRID:SCR_024901) was utilized to test the differential abundances of different genera, species, and pathways between our PD, IBD, and control cohorts.

First, to analyze the differential abundance of different taxa, phyloseq objects were created for genus and species-level taxa respectively. Phyloseq objects were made according to the Phyloseq GitHub tutorial [https://vaulot.github.io/tutorials/Phyloseq_tutorial.html]. Three tables were created using otu_table(), tax_table(), sample_data() functions and combined using the phyloseq() function from the phyloseq R package [https://www.bioconductor.org/packages/release/bioc/html/phyloseq.html] (RRID:SCR_013080). Taxa that were not present in at least 10% of samples were removed. There were 650 genera that were filtered to 140 genera for analysis and 1167 species that were filtered to 233 species for analysis.

Differential abundance was then performed using ancombc2() function from the ANCOMBC R package [https://www.bioconductor.org/packages/release/bioc/html/ANCOMBC.html]. The phyloseq object was used as the input, the fix_formula character string specified how the taxa depended on the fixed effects in the metadata-in this case diagnosis and z-scored total read count were specified, taxa_level was used to specify either species or genus level analysis, p_adj_method was set to BH, group was the name of our cohort in the metadata, lib_cut=0, struc_zero=F, neg_1b=F, alpha = 0.05, global=F, and pairwise=T. Primary output from ANCOM-BC2 contains the log-fold-changes, standard errors, p-values, and adjusted p-values for the examined taxa and specified fixed effects. Pairwise output contains the log-fold-changes, standard errors, p-values, and adjusted p-values for all of the different taxa compared in pairwise analysis between cohorts with control as the reference group. The significant fixed effects in the primary output files and significant taxa emerging from the pairwise comparisons within the pairwise output file were extracted into new data tables. For differential taxonomic abundance data, a genus-level plot was created to visualize the log-fold-change values of different taxa across three different pairwise comparisons of our cohort using ggplot2. To determine if sex, age, and certain medications were associated with differential abundance of any previously identified significant taxa, a multivariate linear regression was performed using the lm() function from that stats package [https://www.rdocumentation.org/packages/stats/versions/3.6.2/topics/lm]. .

For analysis of MetaCyc pathways, a similar approach was taken to create the phyloseq objects, with the difference being MetaCyc pathway information was utilized instead of taxonomic levels and values were RPKs instead of counts. From the pathabundance output files from HUMAnN3.5, all taxonomic information was removed to perform the analysis on a community-level. In the pathway data table, there were 521 distinct pathways, and those that were not present in at least 25% of samples were filtered out. This left 350 pathways for analysis in ANCOM-BC2. The settings used to run ANCOM-BC2 were identical to those used when running the taxonomic differential abundance analysis.

For pathway differential abundance analysis, significant pathways were extracted from the pairwise comparison output files. These pathways were then separated into three separate data frames. Pathways in each data frame were ordered from lowest to highest adjusted p-value (q value) to easily determine the pathways that were found to be most significant in each pairwise comparison performed. Pathways that were found to be significantly associated with PD (from the PD vs control comparison) and IBD (from the IBD vs control comparison) were extracted into a separate shared pathways data frame. A Venn diagram was created to display the PD and IBD associated pathways as well as those that were found to be shared between both conditions using the ggVennDiagram() function from the VennDiagram package.

### Analysis of Larger, Publicly Available Datasets

The open access metagenomic datasets downloaded for this analysis contained available metadata, taxonomic profiling via MetaPhlAn, and functional analysis via HUMAnN. For a larger PD cohort, data from Wallen et al.^23^ was downloaded via Zenodo [https://zenodo.org/doi/10.5281/zenodo.7246184] consisting of 490 PD and 234 Control samples. This dataset is referred to as “Wallen PD” throughout the rest of this paper.

According to their report, the majority of control subjects in this cohort are spousal controls. For a larger IBD cohort, Human Microbiome Project 2 (HMP2) data dated 2018-08-20 was downloaded via [https://www.ibdmdb.org/results]. Due to the large age range of the HMP2 project, this dataset was filtered to include only subjects 40 years and older. This cut off was chosen as it was the age of the youngest subject enrolled in the UFPF study. After age filtering, this dataset consisted of 198 IBD (101 CD, 97 UC) and 139 non-IBD samples. The HMP2 data did not collect spousal or healthy controls, but rather non-IBD subjects refer to subjects that were seen by the GI physician but were not diagnosed with IBD. This dataset is referred to as “HMP2 IBD” throughout the rest of this paper.

Analysis of these datasets generally followed the same methods as outlined above for the UFPF study data. These datasets were analyzed separately due to the numerous differences in collection method, subject geographic location, diet, differences in control groups, etc. Although Wallen et al.^23^ performed many of these analyses in their paper, we have chosen to re-analyze this data with updated tools and stricter parameters. While they utilized ANCOM-BC, the second iteration, ANCOM-BC2, is the updated R package which is more computationally intensive and permits calculation of pairwise comparisons. Additionally, we have maintained an FDR of 0.05 for all analyses. Analysis was performed at the genus and species level. Fixed-effects utilized for the Wallen dataset were kept consistent with those utilized in their publication: case status (diagnosis), collection method, and z-scored total read counts. Fixed-effects for the HMP2 dataset included diagnosis and z-scored total read count. Five HMP2 subjects did not have available functional data, so they were excluded from the functional analysis.

### Relating UFPF Study Data to Larger Datasets

To relate the findings of our UFPF study data with those of the larger, external Wallen and HMP2 datasets, we performed a custom module analysis where we observe the overall abundance of significant features extracted from the Wallen and HMP2 datasets within our UFPF cohorts. Initially, significantly enriched and depleted species were extracted into separate feature lists from the ANCOM-BC2 differential abundance analysis of the Wallen and HMP2 datasets. We then calculated module scores comprised of the average abundance of those species found within those feature lists in a dataset of interest, in this case, within the UFPF PD, IBD, and Control cohorts, using the WGCNA package [https://cran.r-project.org/web/packages/WGCNA/index.html] (RRID:SCR_003302). Quality control check was performed by calculating module scores within in the Wallen and HMP2 datasets, ensuring that the abundance of features matched the directionality seen in the feature lists created initially. The abundance of significant features from the Wallen and HMP2 within the UFPF study data were then plotted. One-way ANOVAs were performed using aov_ez() from the afex package [https://cran.r-project.org/web/packages/afex/index.html] (RRID:SCR_022857). Pairwise comparisons were performed using the emmeans package [https://cran.r-project.org/web/packages/emmeans/index.html] (RRID:SCR_018734) and cld() from the multcomp package [https://cran.r-project.org/web/packages/multcomp/index.html] RRID:SCR_018255) was used to assign letters to denote significance. This module analysis was then performed a second time using significant MetaCyc pathways in the feature lists.

## Supporting information

Key Resource Table

Subject Questionnaires

## Data Availability

The raw, pre-QC metagenomic sequences (with human sequences removed) from the UFPF dataset utilized in this analysis are available to the public with no restrictions at the NCBI Sequence Read Archive (SRA) under BioProject ID PRJNA1096686. The post-QC taxonomic and functional profiling output utilized for downstream analyses outlined in this manuscript are publicly available with no restrictions at Zenodo [https://zenodo.org/doi/10.5281/zenodo.10912505]. The human reference genome used for decontaminating metagenomic sequences (GRCh38.p13) is publicly available at NCBI under the GenBank assembly accession number GCA_000001405.28 [https://www.ncbi.nlm.nih.gov/assembly/GCA_000001405.28]. The ChocoPhlAn database (vJun23) used by MetaPhlAn4 for taxonomic profiling is publicly available for download from the MetaPhlAn databases Segatalab FTP site [http://cmprod1.cibio.unitn.it/biobakery4/metaphlan_databases]. The ChocoPhlAn and UniRef90 reference databases (v201901b) used during functional profiling with HUMAnN3.5 are publicly available to download by using the ‘humann_databases’ utility script as described on the HUMAnN GitHub page [https://github.com/biobakery/humann?tab=readme-ov-file#installation-update]. Functional profiling of pathways by HUMAnN come packaged with the program and were derived from the publicly accessible MetaCyc reference database (v24) [https://metacyc.org/]. All code used to perform sequence QC, taxonomic and functional profiling, statistical analyses, and figure generation are provided to the public with no restrictions on this project’s GitHub page [https://github.com/maevekrueger/UFPF_metagenomics]. A detailed list of the software packages used, RRIDs, links, and version numbers can be found in the Supplementary Materials.

## Acknowledgements

We thank members of the Tansey Lab, Dr. Haydeh Payami at the University of Alabama at Birmingham, Dr. Christian Jobin at the University of Florida, and Dr. Madelyn Houser at Emory University for their helpful critiques. We are grateful to the study participants as well as the movement disorder specialists, GI specialists, and clinical research coordinators for their tireless work recruiting subjects and collecting the samples needed for this study. This project was supported by funds from the Parkinson’s Foundation under the Research Center of Excellence Award to M.G.T (PF-RCE-1945), the Michael J. Fox Foundation and the Aligning Science Across Parkinson’s (ASAP) Collaborative Research Network ‘Circuit’ award to M.G.T and Team Liddle. MJFF administers the grant ASAP-020527 on behalf of ASAP and itself. For the purpose of open access, the author has applied a CC-BY public copyright license to the Author Accepted Manuscript (AAM) version arising from this submission.

## Notes

https://www.ncbi.nlm.nih.gov/bioproject/1096686

https://zenodo.org/doi/10.5281/zenodo.10912505

https://github.com/maevekrueger/UFPF_metagenomics

### Competing Interest Statement

The authors have declared no competing interest.

