## Supplementary material for "A comparative analysis of Parkinson’s disease and inflammatory bowel disease gut microbiomes highlights shared depletions in key butyrate-producing bacteria": Subject Questionnaires

FOR OFFICE USE ONLY

Subject ID: ____________

Please indicate if you or your relatives have any of the following conditions.

| Condition | You | Biological Mother | Biological Father | Biological Brothers or Sisters | Biological Children |
|---|---|---|---|---|---|
| **Neurological** | | | | | |
| Multiple Sclerosis | | | | | |
| Myasthenia Gravis | | | | | |
| Guillain-Barré Syndrome | | | | | |
| CIDP | | | | | |
| Polymyositis | | | | | |
| Dermatomyositis | | | | | |
| Alzheimer's Disease | | | | | |
| Non-Alzhemeier's Dementia | | | | | |
| **Endocrine** | | | | | |
| Hypothyroidism (under-active) | | | | | |
| Hyperthyroidism (over-active) | | | | | |
| Hashimoto thyroiditis | | | | | |
| Grave's disease | | | | | |
| Diabetes mellitus (age started) | | | | | |
| Type 1 | | | | | |
| Type 2 | | | | | |
| Addison's Disease | | | | | |
| Autoimmune Adrenalitis | | | | | |
| **Skin** | | | | | |
| Vitiligo | | | | | |
| Scleroderma | | | | | |
| Pemphigus | | | | | |
| Psoriasis | | | | | |
| **Gastrointestinal** | | | | | |
| Crohn's Disease | | | | | |
| Ulcerative Colitis | | | | | |
| B12 deficiency | | | | | |
| **Bones and Joints** | | | | | |
| Rhematoid Arthritis | | | | | |
| Reiter's Syndrome | | | | | |
| Ankylosing Spondylitis | | | | | |
| **Systemic** | | | | | |
| Lupus (SLE) | | | | | |
| Sjogren's Syndrome | | | | | |
| **Hematologic** | | | | | |
| Hemolytic Anemia | | | | | |
| Aplastic Anemia | | | | | |
| Pernicious Anemia | | | | | |
| Antiphospholipid Syndrome | | | | | |
| Thrombocytopenia | | | | | |
| TTP | | | | | |
| ITP | | | | | |
| Other Autoimmune Disorders | | | | | |
| Cancer | | | | | |
| Specify Type | | | | | |

FOR OFFICE USE ONLY Subject ID:________________

**Medications**

**Name** **Total Daily Dose**

1. ______________________________________________
2. ______________________________________________
3. ______________________________________________
4. ______________________________________________
5. ______________________________________________
6. ______________________________________________
7. ______________________________________________
8. ______________________________________________
9. ______________________________________________
10. ______________________________________________

FOR OFFICE USE ONLY Subject ID:________________

**Tobacco**

Have you smoked at least 100 cigarettes (about 5 packs) in your entire lifetime? ☐Yes ☐No

During the time that you smoked, how much did you smoke on average, and for how many years? Check all that apply.

☐Less than ½ pack per day, for __________ years (*specify number of years*)

☐Equal to or more than ½ pack but less than 1 pack per day, for ___________ years (*specify number of years*)

☐Equal to or more than 1 pack but less than 2 packs per day, for ___________years (*specify number of years*)

☐Equal to or more than 2 packs per day, for ___________years (*specify number of years*)

At what age did you begin smoking? _________________________

Are you still smoking? ☐ Yes ☐ No If not, at what age did you stop? ____________________

**NSAIDs**

***Over the Counter NSAIDs***

NSAIDs are non-steroidal anti-inflammatory drugs, like Ibuprofen, Motrin IB, Advil, and Aleve, which are commonly used for pain. Aspirin and acetaminophen like Tylenol are not NSAIDs.

How often do you (or did you) take **over the counter NSAIDS**, and for how many years? Check all that apply.

☐ Never

☐ Less than once a week, for __________ years (*specify number of years)*

☐ About one to four times a week, for __________ years (*specify number of years)*

☐ About five to ten times a week, for __________ years (*specify number of years)*

☐ More than 10 times a week, for __________ years (*specify number of years)*

FOR OFFICE USE ONLY Subject ID:_______________

***Prescription NSAIDs***

**Prescription NSAIDS** include Anaprox (generic name: naproxen), Arthrotec (diclofenac sodium), Bextra (valdecoxib), Cataflam (diclofenac potassium), Celebrex (celecoxib), Clinoril (sulindac), Dolobid (diflunisal), EC-naprosy n (naproxen), Feldene (piroxicam), Indocin (indomethacin), Mobic (meloxicam), Motrin (ibuprofen), Naprelan (nalproxen controlled release), Naprosyn (naproxen), Ponstel (mefenamic acid), Relafen (nabumetone), Toradol (ketcrolactromethamine), Triisate (choline magnesium salicylate), Vioxx (rofecoxib), Voltaren (diclofenac sodium)

How often do you (or did you) take prescription NSAIDs, and for how many years? Check all that apply.

- ☐ Never
- ☐ Less than once a week, for _________ years (*specify number of years)*
- ☐ About one to four times a week, for _________ years (*specify number of years)*
- ☐ About five to ten times a week, for _________ years (*specify number of years)*
- ☐ More than 10 times a week, for _________ years (*specify number of years)*

**Caffeine**

How much **caffeinated coffee** do you (or did you) drink, and for how many years? Check all that apply.
A cup is about 5 ounces, which is the size of a small Styrofoam cup or a china cup. A coffee mug is 2 cups.

- ☐ Never
- ☐ Less than 2 cups a week, for _________ years (*specify number of years*)
- ☐ 1-2 cups a day, for ___________ years (*specify number of years*)
- ☐ 3-5 cups a day, for ___________ years (*specify number of years*)
- ☐ 6 or more cups a day, for ___________ years (*specify number of years*)

At what age did you start drinking caffeinated coffee? ________

Are you still drinking caffeinated coffee? ☐ Yes ☐ No If not, at what age did you stop? _______________

FOR OFFICE USE ONLY Subject ID:________________

How much **caffeinated tea** do you (or did you) drink, and for how many years? Check all that apply.
A cup is about 5 ounces, which is the size of a small Styrofoam cup or a china cup. A coffee mug is 2 cups.

- ☐ Never
- ☐ Less than 2 cups a week, for __________ years (*specify number of years*)
- ☐ 1-2 cups a day, for ____________ years (*specify number of years*)
- ☐ 3-5 cups a day, for ____________ years (*specify number of years*)
- ☐ 6 or more cups a day, for ____________ years (*specify number of years*)

At what age did you start drinking caffeinated tea? _________
Are you still drinking caffeinated tea? ☐ Yes ☐ No If not, at what age did you stop? ________________

How much **caffeinated soda** do you (or did you) drink, and for how many years? Check all that apply.
A can of soda is 12 oz.

- ☐ Never
- ☐ Less than 2 cups a week, for __________ years (*specify number of years*)
- ☐ 1-2 cups a day, for ____________ years (*specify number of years*)
- ☐ 3-5 cups a day, for ____________ years (*specify number of years*)
- ☐ 6 or more cups a day, for ____________ years (*specify number of years*)

At what age did you start drinking caffeinated soda? _________
Are you still drinking caffeinated soda? ☐ Yes ☐ No If not, at what age did you stop? ________________

**Head Injury**

Have you ever had a head injury that caused loss of consciousness or required medical care? ☐ Yes ☐ No (If no, go to Ethnic origins section)
How many times in your life have you had such a head injury? _______ times
How old were you when you had your first head injury? _________ years old
How old were you when you had your last head injury? _________ years old

FOR OFFICE USE ONLY        Subject ID:______________

**Ethnic Origins**

From what countries did your paternal ancestors immigrate to the United States? ______________________________

From what countries did your maternal ancestors immigrate to the United States? ______________________________

Are you Hispanic or Latino? ☐ Yes ☐ No

What race do you most identify yourself with:

☐ American Indian/Alaskan Native

☐ Asian

☐ Native Hawaiian or other Pacific Islander

☐ Black or African American

☐ White

☐ More than one race

If you are from a particular religious lineage, such as Jewish, Amish, etc., please specify.
This is a question of genetic lineage, not personal preference: ______________________________

**Immune System Related Events**

Any recent immunizations (including annual flu)? Please provide type and date.

Have you had a fever or a known infection in the past 30 days? Please specify.

Have you been hospitalized in the past 30 days? Please specify.

Have you had surgery in the past 30 days? Please specify.

### Activities of Daily Living Scale

Patient: ____________ Date: ____________ Time Point: B 2/3w 3m 1y

Your Name: ____________ Relationship to Participant: ____________

Please circle the description under each heading which best characterizes the patient's behavior over the *past several weeks*.

**Instrumental Activities Scale**

Ability to Use the Telephone

(1) Operates telephone on own initiative, looks up and dials numbers, etc.
(2) Answers telephone; dials a few well-known numbers, but does not look up or dial less-frequently used numbers without assistance.
(3) Does not use telephone at all.

Shopping

(1) Takes care of all shopping needs independently (i.e. goes to the store, selects needed items, pays for items and brings them home).
(2) Shops independently for small purchases.
(3) Completely unable to shop alone.

Food Preparation

(1) Plans, prepares and serves adequate meals independently; manages the stove without help.
(2) Prepares adequate meals if supplied with ingredients; heats, serves and prepares meals but does not maintain an adequate diet.
(3) Needs to have meals prepared and served.

Housekeeping

(1) Maintains house (house-cleaning, vacuum-cleaning, washes floors, etc.) alone or with occasional assistance
(2) Performs light daily tasks such as dish-washing and bed-making, but cannot maintain an acceptable level of cleanliness.
(3) Does not participate in any housekeeping tasks.

Laundry

(1) Does personal laundry completely.
(2) Launders small items, washes some clothes by hand, needs help with major laundry.
(3) All laundry must be done by others.

Mode of Transportation

(1) Travels independently on public transportation, drives own car, or arranges own travel via taxi.
(2) Travels on public transportation, taxi, or automobile when assisted or accompanied by another.
(3) Does not travel at all.

Medications

(1) Is responsible for taking medication in correct dosages at correct time
(2) Takes responsibility of medication if it is prepared in advance in separate dosage.
(3) Is not capable of dispensing own medication.

Ability to Handle Finances

(1) Manages financial matters independently (budgets, writes checks, pays rent, bills, collects and keeps track of income, etc.).
(2) Manages day-to-day purchases but needs help with banking, major purchases, etc.
(3) Incapable of handling money.

**Physical Self Maintenance Scale**

Toilet

(1) Cares for self at toilet completely, no incontinence
(2) Needs to be reminded, needs help cleaning self or redressing, or has rare (weekly at most) accidents.
(3) Soiling of wetting more than once a week or total incontinence.

Feeding

(1) Eats without assistance.
(2) East with mild to moderate assistance at mealtimes and/or with special preparation of food or help in cleaning up.
(3) Does not feel self at all

Dressing

(1) Dresses, undresses and selects clothes from wardrobe without help.
(2) Needs minor to moderate assistance in dressing and/or in the selection of clothes.
(3) Unable to dress self.

Hair Grooming

(1) Washes and combs/brushes hair independently.
(2) Needs moderate and regular assistance in grooming or does not wash/comb hair unless told.
(3) Caregiver must wash and comb hair.

Dental Hygiene

(1) Brushes teeth independently and does not require to be reminded.
(2) Needs moderate and regular supervision or assistance in brushing teeth (needs to be reminded to brush teeth; needs help gathering all necessary objects, etc.)
(3) Can not brush teeth without complete assistance.

Nail Care

(1) Cares for nails (cleans and clips) regularly and without assistance.
(2) Must be reminded to care for nails or requires some assistance in cutting nails.
(3) Relies on others for total nail grooming care.

Bathing

(1) Bathes self (tub, shower, sponge bath) without help.
(2) Needs supervision or assistance in bathing.
(3) Does not wash self.

### BDI - II

Instructions: This questionnaire consists of 21 groups of statements. Please read each group of statements carefully. And then pick out the one statement in each group that best describes the way you have been feeling during the past two weeks, including today. Circle the number beside the statement you have picked. If several statements in the group seem to apply equally well, circle the highest number for that group. Be sure that you do not choose more than one statement for any group, including Item 16 (Changes in Sleeping Pattern) or Item 18 (Changes in Appetite).

1. Sadness
   0. I do not feel sad.
   1. I feel sad much of the time.
   2. I am sad all the time.
   3. I am so sad or unhappy that I can't stand it.

2. Pessimism
   0. I am not discouraged about my future.
   1. I feel more discouraged about my future than I used to.
   2. I do not expect things to work out for me.
   3. I feel my future is hopeless and will only get worse.

3. Past Failure
   0. I do not feel like a failure.
   1. I have failed more than I should have.
   2. As I look back, I see a lot of failures.
   3. I feel I am a total failure as a person.

4. Loss of Pleasure
   0. I get as much pleasure as I ever did from the things I enjoy.
   1. I don't enjoy things as much as I used to.
   2. I get very little pleasure from the things I used to enjoy.
   3. I can't get any pleasure from the things I used to enjoy.

5. Guilty Feelings
   0. I don't feel particularly guilty.
   1. I feel guilty over many things I have done or should have done.
   2. I feel quite guilty most of the time.
   3. I feel guilty all of the time.

6. Punishment Feelings
   0. I don't feel I am being punished.
   1. I feel I may be punished.
   2. I expect to be punished.
   3. I feel I am being punished.

7. Self-Dislike
   0. I feel the same about myself as ever.
   1. I have lost confidence in myself.
   2. I am disappointed in myself.
   3. I dislike myself.

8. Self-Criticalness

0. I don't criticize or blame myself more than usual.
1. I am more critical of myself than I used to be.
2. I criticize myself for all of my faults.
3. I blame myself for everything bad that happens.

9. Suicidal Thoughts or Wishes

0. I don't have any thoughts of killing myself.
1. I have thoughts of killing myself, but I would not carry them out.
2. I would like to kill myself.
3. I would kill myself if I had the chance.

10. Crying

0. I don't cry anymore than I used to.
1. I cry more than I used to.
2. I cry over every little thing.
3. I feel like crying, but I can't.

11. Agitation

0. I am no more restless or wound up than usual.
1. I feel more restless or wound up than usual.
2. I am so restless or agitated, it's hard to stay still.
3. I am so restless or agitated that I have to keep moving or doing something.

12. Loss of Interest

0. I have not lost interest in other people or activities.
1. I am less interested in other people or things than before.
2. I have lost most of my interest in other people or things.
3. It's hard to get interested in anything.

13. Indecisiveness

0. I make decisions about as well as ever.
1. I find it more difficult to make decisions than usual.
2. I have much greater difficulty in making decisions than I used to.
3. I have trouble making any decisions.

14. Worthlessness

0. I do not feel I am worthless.
1. I don't consider myself as worthwhile and useful as I used to.
2. I feel more worthless as compared to others.
3. I feel utterly worthless.

15. Loss of Energy

0. I have as much energy as ever.
1. I have less energy than I used to have.
2. I don't have enough energy to do very much.
3. I don't have enough energy to do anything.

16. Changes in Sleeping Pattern

0. I have not experienced any change in my sleeping.
1a I sleep somewhat more than usual.
1b I sleep somewhat less than usual.
2a I sleep a lot more than usual.
2b I sleep a lot less than usual.
3a I sleep most of the day.
3b I wake up 1-2 hours early and can't get back to sleep.

17. Irritability

0. I am not more irritable than usual.
1. I am more irritable than usual.
2. I am much more irritable than usual.
3. I am irritable all the time.

18. Changes in Appetite

0. I have not experienced any change in my appetite.
1a My appetite is somewhat less than usual.
1b My appetite is somewhat greater than usual.
2a My appetite is much less than before.
2b My appetite is much greater than usual.
3a I have no appetite at all.
3b I crave food all the time.

19. Concentration Difficulty

0. I can concentrate as well as ever.
1. I can't concentrate as well as usual.
2. It's hard to keep my mind on anything for very long.
3. I find I can't concentrate on anything.

20. Tiredness or Fatigue

0. I am no more tired or fatigued than usual.
1. I get more tired or fatigued more easily than usual.
2. I am too tired or fatigued to do a lot of the things I used to do.
3. I am too tired or fatigued to do most of the things I used to do.

21. Loss of Interest in Sex

0. I have not noticed any recent change in my interest in sex.
1. I am less interested in sex than I used to be.
2. I am much less interested in sex now.
3. I have lost interest in sex completely.

Total Score: _______

THE PSYCHOLOGICAL CORPORATION
Harcourt Brace & Company
Copyright 1996, by Aaron T. Beck. All rights reserved.

**PDBP**
Parkinson's Disease
Biomarkers Program

* Required fields

### Modified Schwab and England Activities of Daily Living Scale

| | |
|---|---|
| *Name of Site: ____________ | *Type of Visit: ____________<br>e.g. Screening, Baseline, 6 months, 12 months, 18 months, 24 months, 30 months, 36 months, 42 months, 48 months, 54 months, 60 months. |
| *Date of Visit: ____________ | *GUID: ____________ |
| *Age of Subject (years and months): ________ | Subject ID: ____________ |

*Score: __________ (Number between 1-100)

100% – Completely independent. Able to do all chores without slowness, difficulty or impairment. Essentially normal. Unaware of any difficulty.

90% – Completely independent. Able to do all chores with some degree of slowness, difficulty and impairment. Might take twice as long. Beginning to be aware of difficulty.

80% – Completely independent in most chores. Takes twice as long. Conscious of difficulty and slowness.

70% – Not completely independent. More difficulty with some chores. Three to four times as long in some. Must spend a large part of the day with chores.

60% – Some dependency. Can do most chores, but exceedingly slowly and with much effort. Errors; some impossible.

50% – More dependent. Help with half, slower, et cetera. Difficulty with everything.

40% – Very dependent. Can assist with all chores, but few alone.

30% – With effort, now and then does a few chores alone or begins alone. Much help needed.

20% – Nothing alone. Can be a slight help with some chores. Severe invalid.

10% – Total dependent, helpless. Complete invalid.

0% – Vegetative functions such as swallowing, bladder and bowel functions are not functioning. Bed-ridden.

NAME :
Education :                Date of birth :
Sex :                DATE :

### MONTREAL COGNITIVE ASSESSMENT (MOCA)
Version 7.1 Original Version

#### VISUOSPATIAL / EXECUTIVE

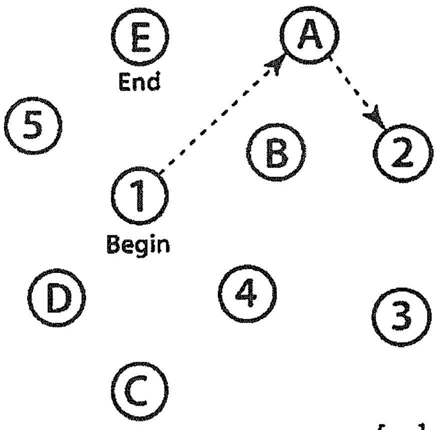


[ ]

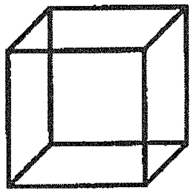

Copy cube

[ ]

Draw CLOCK (Ten past eleven)
(3 points)

[ ] Contour    [ ] Numbers    [ ] Hands

POINTS: ___/5

#### NAMING

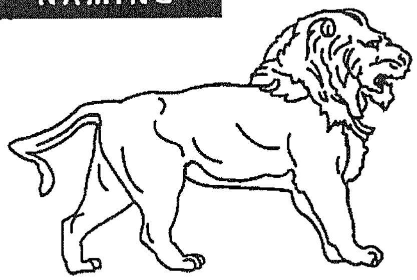

[ ]

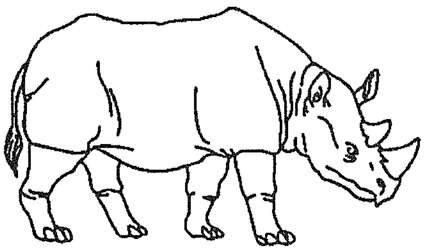

[ ]

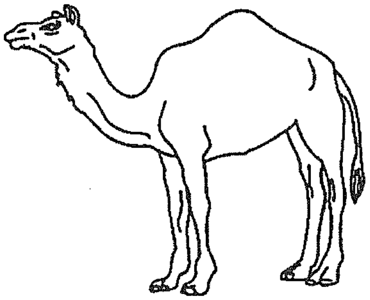

[ ]

___/3

#### MEMORY

Read list of words, subject must repeat them. Do 2 trials, even if 1st trial is successful. Do a recall after 5 minutes.

| | FACE | VELVET | CHURCH | DAISY | RED |
|---|---|---|---|---|---|
| 1st trial | | | | | |
| 2nd trial | | | | | |

No points

#### ATTENTION

Read list of digits (1 digit/ sec.). Subject has to repeat them in the forward order [ ] 2 1 8 5 4
Subject has to repeat them in the backward order [ ] 7 4 2
___/2

Read list of letters. The subject must tap with his hand at each letter A. No points if ≥ 2 errors
[ ] F B A C M N A A J K L B A F A K D E A A A J A M O F A A B
___/1

Serial 7 subtraction starting at 100 [ ] 93 [ ] 86 [ ] 79 [ ] 72 [ ] 65
4 or 5 correct subtractions: **3 pts**, 2 or 3 correct: **2 pts**, 1 correct: **1 pt**, 0 correct: **0 pt**
___/3

#### LANGUAGE

Repeat : I only know that John is the one to help today. [ ]
The cat always hid under the couch when dogs were in the room. [ ]
___/2

Fluency / Name maximum number of words in one minute that begin with the letter F [ ] _____ (N ≥ 11 words)
___/1

#### ABSTRACTION

Similarity between e.g. banana - orange = fruit [ ] train – bicycle [ ] watch - ruler
___/2

#### DELAYED RECALL

| | FACE | VELVET | CHURCH | DAISY | RED | |
|---|---|---|---|---|---|---|
| Has to recall words WITH NO CUE | [ ] | [ ] | [ ] | [ ] | [ ] | Points for UNCUED recall only |
| Optional: Category cue | | | | | | |
| Multiple choice cue | | | | | | |

___/5

#### ORIENTATION

[ ] Date [ ] Month [ ] Year [ ] Day [ ] Place [ ] City
___/6

© Z.Nasreddine MD **www.mocatest.org** Normal ≥26 / 30

TOTAL ___/30
Add 1 point if ≤ 12 yr edu

Administered by: ________________

FOR OFFICE USE ONLY Subject ID:_______________

### Confidential Environmental and Family History Questionnaire for PD Studies

Today's Date: ________________________

**Personal Information**

Name: ________________________ Birthdate: ________________ Sex: M☐ F☐

Address: ________________________________________________

Phone (H): ________________ Phone (W): ________________ Email: ________________

Have you ever been diagnosed with, or suspected to have Parkinson's disease or parkinsonism? ☐ Yes ☐ No

If you answered **yes** to the above question:

What was your first symptom? ________________

At what age did you notice the first symptom? __________

If seen by a physician, at what age were you diagnosed? ______ Name of Neurologist (City, State):________________

If this form is completed by someone other than the subject/patient:

Name of person completing form: ________________________________

Relationship to subject/patient: ________________ Number of years you have known the subject/patient: __________

Address: ________________________________________________

Phone (H): ________________ Phone (W): ________________ Email: ________________

**Parkinson's Disease Screening Questionnaire (Control)**



Site Name: ____________________

Subject ID: ______________

1. Do you have trouble arising from a chair? ☐YES ☐NO
2. Is your handwriting smaller than it once was? ☐YES ☐NO
3. Do people tell you that your voice is softer than it once was? ☐YES ☐NO
4. Is your balance poor? ☐YES ☐NO
5. Do your feet ever seem to get stuck to the floor? ☐YES ☐NO
6. Do people tell you that your face seems less expressive than it once did? ☐YES ☐NO
7. Do your arms and legs shake? ☐YES ☐NO
8. Do you have trouble buttoning buttons? ☐YES ☐NO
9. Do you shuffle your feet and / or take tiny steps when you walk? ☐YES ☐NO
10. Has anyone ever told you that you have Parkinson's disease? ☐YES ☐NO
11. Have you ever taken levodopa or Sinemet? ☐YES ☐NO

---

Rocca et al., J Clin Epidemiol Vol. 51, No. 6, pp. 517 – 523, 1998

### Unified PARKINSON Disease Rating Scale (UPDRS)

(AAN, 1995)

*The UPDRS is a rating tool to follow the longitudinal course of Parkinson's Disease. It is made up of the 1)Mentation, Behavior, and Mood, 2)ADL and 3)Motor sections. These are evaluated by interview. Some sections require multiple grades assigned to each extremity.*

**I. Mentation, Behavior, Mood**

o *Intellectual Impairment*
- 0-none
- 1-mild (consistent forgetfulness with partial recollection of events with no other difficulties)
- 2-moderate memory loss with disorientation and moderate difficulty handling complex problems
- 3-severe memory loss with disorientation to time and often place, severe impairment with problems
- 4-severe memory loss with orientation only to person, unable to make judgments or solve problems

o *Thought Disorder*
- 0-none
- 1-vivid dreaming
- 2-"benign" hallucination with insight retained
- 3-occasional to frequent hallucination or delusions without insight, could interfere with daily activities
- 4-persistent hallucination, delusions, or florid psychosis.

o *Depression*
- 0-not present
- 1-periods of sadness or guilt greater than normal, never sustained for more than a few days or a week
- 2-sustained depression for >1 week
- 3-vegetative symptoms (insomnia, anorexia, abulia, weight loss)
- 4-vegetative symptoms with suicidality

o *Motivation/Initiative*
- 0-normal
- 1-less of assertive, more passive
- 2-loss of initiative or disinterest in elective activities
- 3-loss of initiative or disinterest in day to say (routine) activities
- 4-withdrawn, complete loss of motivation

**II. Activities of Daily Living**

o *Speech*
- 0-normal
- 1-mildly affected, no difficulty being understood
- 2-moderately affected, may be asked to repeat
- 3-severely affected, frequently asked to repeat
- 4-unintelligible most of time

o *Salivation*
- 0-normal
- 1-slight but noticeable increase, may have nighttime drooling
- 2-moderately excessive saliva, hay minimal drooling
- 3-marked drooling

o *Swallowing*
- 0-normal
- 1-rare choking
- 2-occasional choking
- 3-requires soft food
- 4-requires NG tube or G-tube

o *Handwriting*
- 0-normal
- 1-slightly small or slow
- 2-all words small but legible
- 3-severely affected, not all words legible
- 4-majority illegible

o *Cutting Food/Handing Utensils*
- 0-normal
- 1-somewhat slow and clumsy but no help needed
- 2-can cut most foods, some help needed
- 3-food must be cut, but can feed self
- 4-needs to be fed

o *Dressing*
- 0-normal
- 1-somewhat slow, no help needed
- 2-occasional help with buttons or arms in sleeves
- 3-considerable help required but can do something alone
- 4-helpless

o *Hygiene*
- 0-normal
- 1-somewhat slow but no help needed
- 2-needs help with shower or bath or very slow in hygienic care
- 3-requires assistance for washing, brushing teeth, going to bathroom
- 4-helpless

o *Turning in Bed/ Adjusting Bed Clothes*
- 0-normal
- 1-somewhat slow no help needed
- 2-can turn alone or adjust sheets but with great difficulty
- 3-san initiate but not turn or adjust alone
- 4-helpless

- o *Falling-Unrelated to Freezing*
  - 0-none
  - 1-rare falls
  - 2-occasiona, less than one per day
  - 3-average of once per day
  - 4->1 per day

- o *Freezing When Walking*
  - 0-normal
  - 1-rare, may have start hesitation
  - 2-occasional falls from freezing,
  - 3-frequent freezing, occasional falls
  - 4-frequent falls from freezin

- o *Walking*
  - 0-normal
  - 1-mild difficulty, day drag legs or decrease arm swing
  - 2-moderate difficultly requires no assist
  - 3-severe disturbance requires assistance
  - 4-cannot walk at all even with assist

- o *Tremor*
  - 0-absent
  - 1-slight and infrequent, not bothersome to patient
  - 2-moderate, bothersome to patient
  - 3-severe, interfere with many activities
  - 4-marked, interferes with many activities

- o *Sensory Complaints Related to Parkinsonism*
  - 0-none
  - 1-occasionally has numbness, tingling, and mild aching
  - 2-frequent, but not distressing
  - 3-frequent painful sensation
  - 4-excruciating pain

**III. Motor Exam**

- o *Speech*
  - 0-normal
  - 1-slight loss of expression, diction,volume
  - 2-monotone, slurred but understandable, mod. impaired
  - 3-marked impairment, difficult to understand
  - 4-unintelligible

- o Facial Expression
  - 0-Normal
  - 1-slight hypomymia, could be poker face
  - 2-slight but definite abnormal diminution in expression
  - 3-mod. hypomimia, lips parted some of time
  - 4-masked or fixed face, lips parted 1/4 of inch or more with complete loss of expression

- o *Tremor at Rest*
  - + Face
    - 0-absent
    - 1-slight and infrequent
    - 2-mild and present most of time
    - 3-moderate and present most of time
    - 4-marked and present most of time
  - + Right Upper Extremity (RUE)
    - 0-absent
    - 1-slight and infrequent
    - 2-mild and present most of time
    - 3-moderate and present most of time
    - 4-marked and present most of time
  - + LUE
    - 0-absent
    - 1-slight and infrequent
    - 2-mild and present most of time
    - 3-moderate and present most of time
    - 4-marked and present most of time
  - + RLE
    - 0-absent
    - 1-slight and infrequent
    - 2-mild and present most of time
    - 3-moderate and present most of time
    - 4-marked and present most of time
  - + LLE
    - 0-absent
    - 1-slight and infrequent
    - 2-mild and present most of time
    - 3-moderate and present most of time
    - 4-marked and present most of time

- o *Action or Postural Tremor*
  - + RUE
    - 0-absent
    - 1-slight, present with action
    - 2-moderate, present with action
    - 3-moderate present with action and posture holding
    - 4-marked, interferes with feeding
  - + LUE
    - 0-absent
    - 1-slight, present with action
    - 2-moderate, present with action
    - 3-moderate present with action and posture holding
    - 4-marked, interferes with feeding

- o *Rigidity*
  - + Neck
    - 0-absent
    - 1-slight or only with activation
    - 2-mild/moderate
    - 3-marked, full range of motion
    - 4-severe
  - + RUE
    - 0-absent
    - 1-slight or only with activation
    - 2-mild/moderate
    - 3-marked, full range of motion
    - 4-severe
  - + LUE
    - 0-absent
    - 1-slight or only with activation
    - 2-mild/moderate
    - 3-marked, full range of motion
    - 4-severe
  - + RLE
    - 0-absent
    - 1-slight or only with activation
    - 2-mild/moderate
    - 3-marked, full range of motion
    - 4-severe
  - + LLE
    - 0-absent

1-slight or only with activation
2-mild/moderate
3-marked, full range of motion
4-severe

- *Finger taps*
  - + Right
    0-normal
    1-mild slowing, and/or reduction in amp.
    2-moderate impaired. Definite and early fatiguing, may have occasional arrests
    3-severely impaired. Frequent hesitations and arrests.
    4-can barely perform
  - + Left
    0-normal
    1-mild slowing, and/or reduction in amp.
    2-moderate impaired. Definite and early fatiguing, may have occasional arrests
    3-severely impaired. Frequent hesitations and arrests.
    4-can barely perform

- *Hand Movements (open and close hands in rapid succession)*
  - + Right
    0-normal
    1-mild slowing, and/or reduction in amp.
    2-moderate impaired. Definite and early fatiguing, may have occasional arrests
    3-severely impaired. Frequent hesitations and arrests.
    4-can barely perform
  - + Left
    0-normal
    1-mild slowing, and/or reduction in amp.
    2-moderate impaired. Definite and early fatiguing, may have occasional arrests
    3-severely impaired. Frequent hesitations and arrests.
    4-can barely perform

- *Rapid Alternating Movements (pronate and supinate hands)*
  - + Right
    0-normal
    1-mild slowing, and/or reduction in amp.
    2-moderate impaired. Definite and early fatiguing, may have occasional arrests
    3-severely impaired. Frequent hesitations and arrests.
    4-can barely perform
  - + Left
    0-normal
    1-mild slowing, and/or reduction in amp.
    2-moderate impaired. Definite and early fatiguing, may have occasional arrests
    3-severely impaired. Frequent hesitations and arrests.
    4-can barely perform

- *Leg Agility (tap heel on ground, amp should be 3 inches)*
  - + Right
    0-normal
    1-mild slowing, and/or reduction in amp.
    2-moderate impaired. Definite and early fatiguing, may have occasional arrests
    3-severely impaired. Frequent hesitations and arrests.
    4-can barely perform
  - + Left
    0-normal
    1-mild slowing, and/or reduction in amp.
    2-moderate impaired. Definite and early fatiguing, may have occasional arrests
    3-severely impaired. Frequent hesitations and arrests.
    4-can barely perform

- *Arising From Chair (pt. arises with arms folded across chest)*
  0-normal
  1-slow, may need more than one attempt
  2-pushes self up from arms or seat
  3-tends to fall back, may need multiple tries but can arise without assistance
  4-unable to arise without help

- *Posture*
  0-normal erect
  1-slightly stooped, could be normal for older person
  2-definitely abnormal, mod. stooped, may lean to one side
  3-severely stooped with kyphosis
  4-marked flexion with extreme abnormality of posture

- *Gait*
  0-normal
  1-walks slowly, may shuffle with short steps, no festination or propulsion
  2-walks with difficulty, little or no assistance, some festination, short steps or propulsion
  3-severe disturbance, frequent assistance
  4-cannot walk

- *Postural Stability (retropulsion test)*
  0-normal
  1-recovers unaided
  2-would fall if not caught
  3-falls spontaneously
  4-unable to stand

- *Body Bradykinesia/ Hypokinesia*
  0-none
  1-minimal slowness, could be normal, deliberate character
  2-mild slowness and poverty of movement, definitely abnormal, or dec. amp. of movement
  3-moderate slowness, poverty, or small amplitude
  4-marked slowness, poverty, or amplitude
