## Supplementary Figures for "A comparative analysis of Parkinson’s disease and inflammatory bowel disease gut microbiomes highlights shared depletions in key butyrate-producing bacteria"

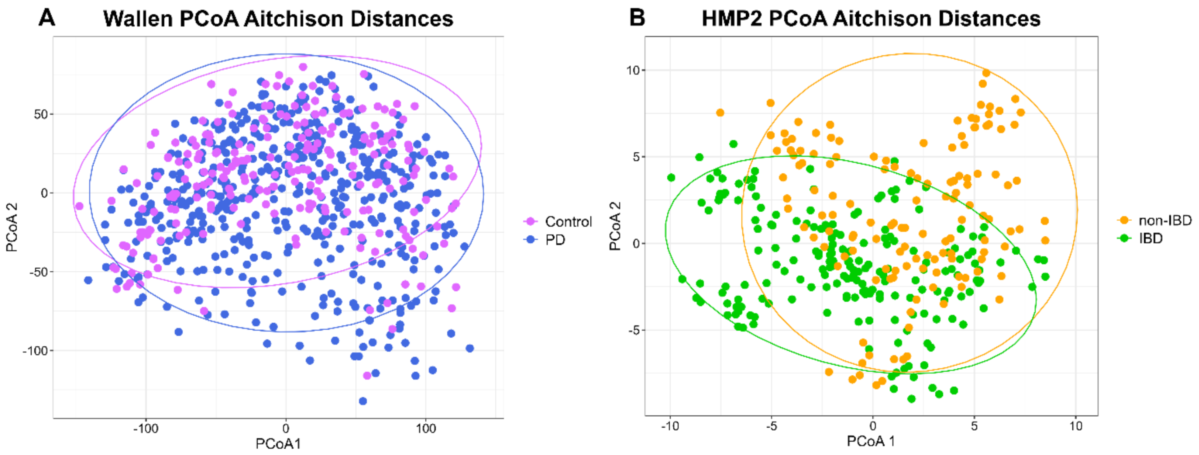


**Supplementary Figure 1. Wallen PD and HMP2 IBD PCoA.** (A) Principal coordinate analysis (PCoA) using Aitchison distances between 490 PD (blue) and 234 neurologically healthy control (pink) metagenomes with all species demonstrated PD dispersion was significantly different than Control by PERMANOVA (*p*=0.001) and PERMDISP (*p*=0.001). (B) PCoA using Aitchison distances between 198 IBD and 139 nonIBD metagenomes with all species demonstrated IBD dispersion was significantly different than nonIBD by PERMANOVA (*p*=0.001) but not by PERMDISP (*p*=0.832).

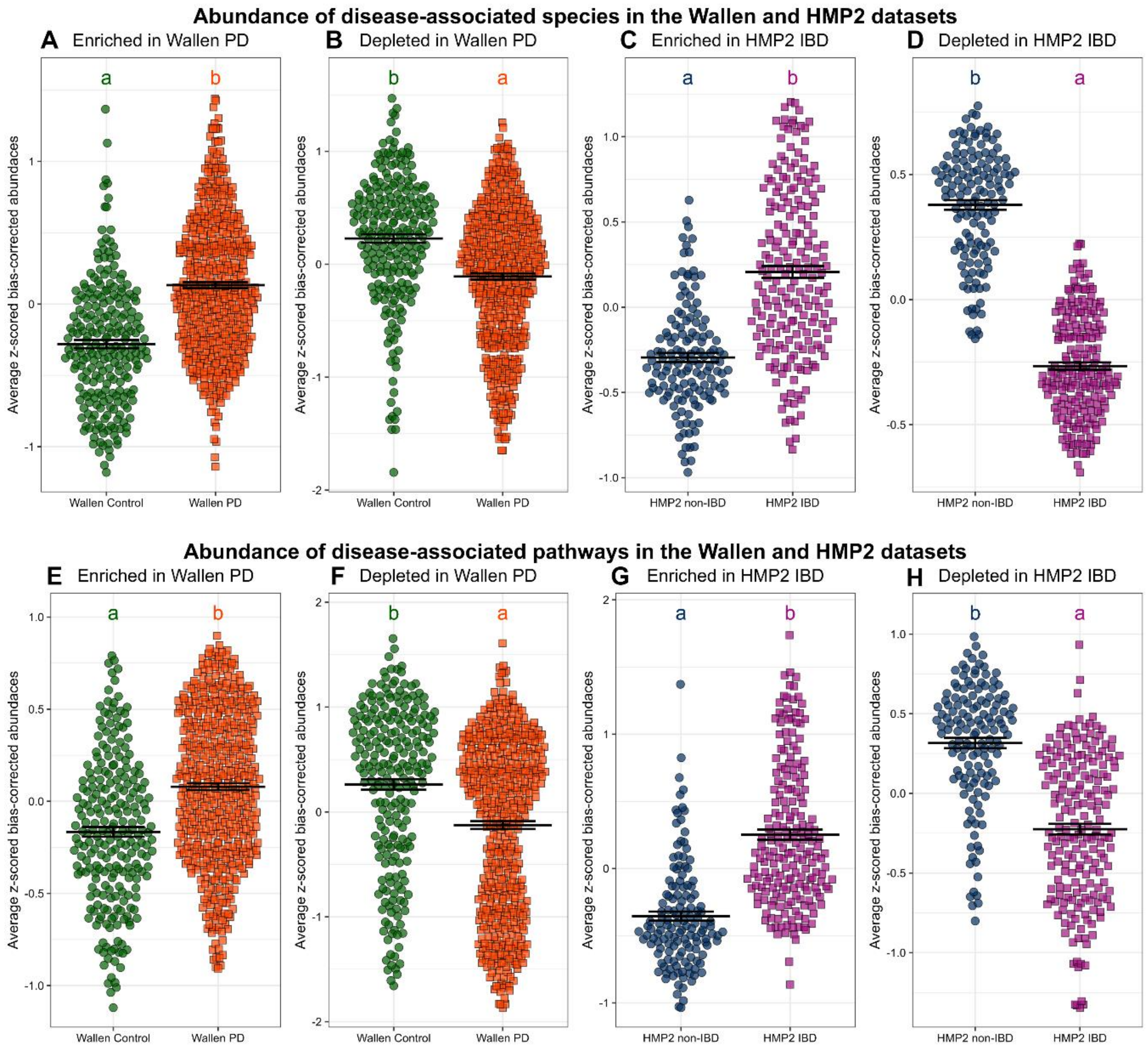


**Supplementary Figure 2. Species and pathway module analysis quality control comparing Wallen PD and HMP2 IBD to their own modules**. Quality control was performed by calculating module scores within in the Wallen and HMP2 datasets, ensuring that the abundance of features match the movement seen in the feature lists created initially. (A-B) The abundance of significant species from the Wallen PD dataset within the Wallen PD cohorts are plotted. (C-D) The abundance of significant species from the HMP2 IBD dataset within the HMP2 IBD cohorts are plotted. Modules are denoted by the title on each individual plot. Significantly enriched and depleted species were extracted into separate feature lists from the ANCOM-BC2 differential abundance analysis of the Wallen and HMP2 datasets. Calculated module scores were comprised of the average abundance of those species found within the Wallen and HMP2 datasets respectively, using the R WGCNA package. One-way ANOVAs were performed followed by calculation of estimated marginal means and pairwise comparisons. Compact letter display was used to display pairwise comparisons, as letters that are different from each other are significantly different ($p < 0.05$). (E-F) The abundance of significant MetaCyc pathways from the Wallen PD dataset within the Wallen PD cohorts are plotted. (G-H) The abundance of significant MetaCyc pathways from the HMP2 IBD dataset within the HMP2 IBD cohorts are plotted. Statistical methods were the same as described above, but the feature lists were comprised of significantly enriched and depleted MetaCyc pathways from the

ANCOM-BC2 differential abundance analysis of the Wallen and HMP2 datasets, and calculated module scores were comprised of the average abundance of those pathways found within those feature lists in the Wallen and HMP2 datasets respectively.
